# CARIBOU: Computational AI Research Interface for Bioinformatics, Omics, and Unifying Agents

**DOI:** 10.64898/2026.05.25.727730

**Authors:** Dylan Riffle, Nima Shirooni, Pavan Sureshkumar, Varshini Vijay, Matthew F. Rose

## Abstract

The growing gap between biological data generation and the availability of expert analysts motivates the development of AI systems capable of autonomously performing meaningful computational biology workflows. Here, we present CARIBOU (Computational AI Research Interface for Bioinformatics, Omics, and Unifying Agents), a multi-agent framework designed for practical deployment within institutional research computing environments. CARIBOU organizes specialized AI agents through researcher-modifiable blueprints that encode analytical roles, domain knowledge, and workflow guidance. All analyses are executed within reproducible computational environments compatible with Singularity/Apptainer-based high-performance computing (HPC) systems commonly used in academic research, while maintaining a persistent shared analytical state across multiple stages of analysis. This design enables CARIBOU to iteratively execute, troubleshoot, and refine bioinformatics workflows rather than simply generate static code. We evaluate CARIBOU across unit-task benchmarks, a metadata reconstruction challenge spanning six public datasets, and two end-to-end single-cell RNA-seq analyses using Allen Brain Atlas hippocampus and Tabula Sapiens large intestine datasets, alongside qualitative case studies demonstrating adaptive reasoning during analysis execution.

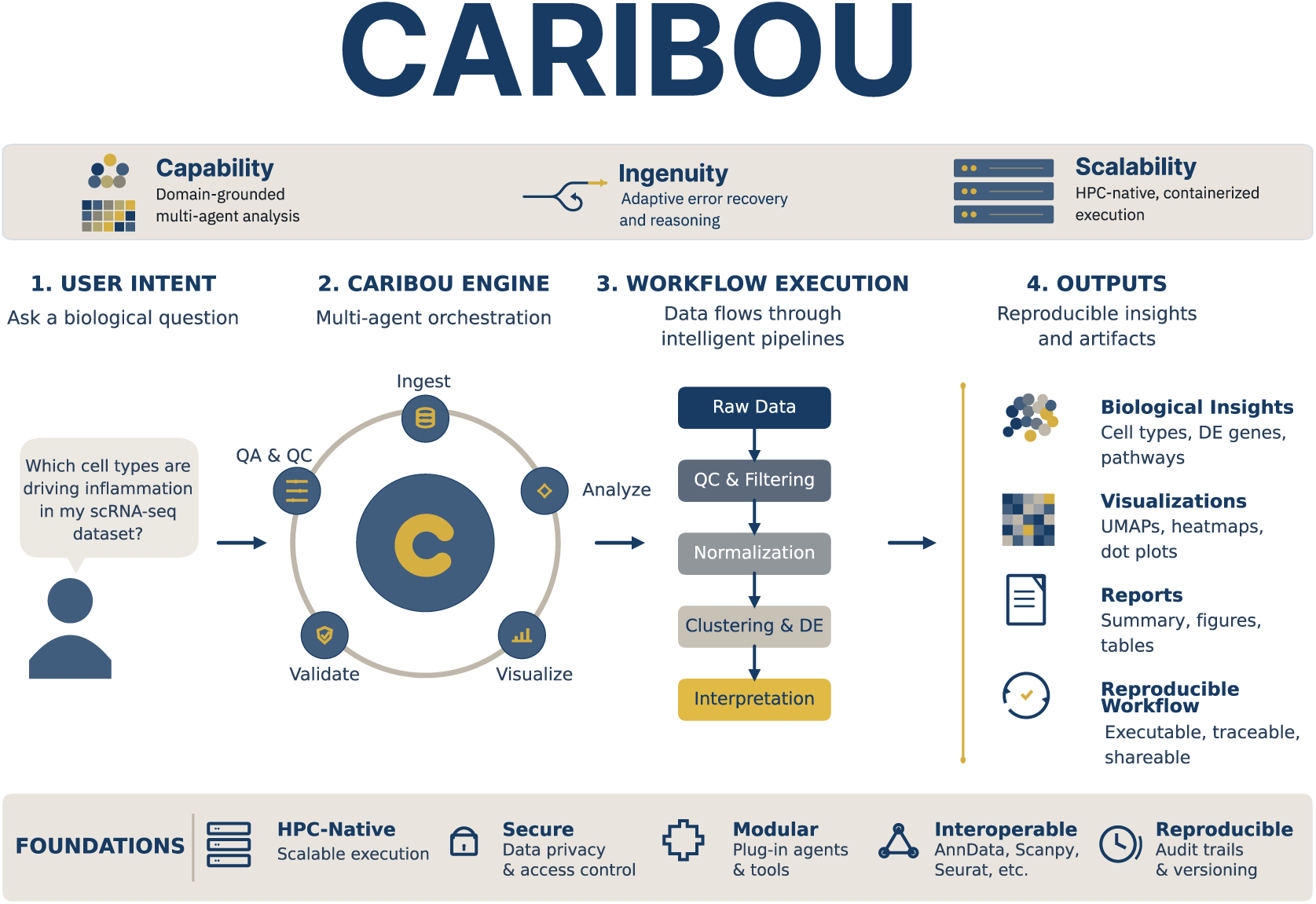

**IN BRIEF:** Modern single-cell and spatial omics studies generate datasets that are increasingly difficult to analyze manually, creating a growing bottleneck between data generation and biological discovery. CARIBOU is a multi-agent AI system designed to autonomously perform bioinformatics analyses within the high-performance computing environments used by research institutions. By combining specialized AI agents with persistent execution environments and built-in analytical workflows, CARIBOU can adaptively execute, troubleshoot, and document complex analyses while maintaining reproducibility and compatibility with real-world research infrastructure.

## INTRODUCTION

Three compounding advances define the present moment in bioscience. The capacity to generate biological data has undergone a step change: single-cell RNA sequencing (scRNA-seq) has moved from a specialized technique to a near-routine assay, and public repositories such as the Gene Expression Omnibus (GEO) (Clough et al., 2024) and the CZI CELLxGENE Census now collectively catalog thousands of single-cell datasets spanning diverse organisms, tissues, and disease states. Computational infrastructure has expanded in parallel; institutional high performance computing (HPC) clusters are now standard in research universities, offering thousands of cores alongside substantial memory and storage. Third, the analytical tooling ecosystem has diversified substantially, from foundational frameworks like Scanpy (Wolf et al., 2018) and Seurat (Hao et al., 2021) to the broad landscape of specialized methods now covering every analytical decision from quality control to differential expression (Luecken and Theis, 2019).

The compound effect of these advances is a dataset commons that is structurally underanalyzed. Most published datasets are analyzed once at the time of publication, using methods that become outdated and pipelines that may not be reproducible (Luecken and Theis, 2019). The bottleneck is not hardware. HPC clusters sit underutilized not because they lack capacity, but because meaningful single-cell analysis is not a matter of applying a fixed computational procedure. It requires a sequence of adaptive decisions: assessing data quality and setting thresholds appropriate to the specific dataset, characterizing batch effects and selecting a correction strategy suited to their structure, validating clustering against biological expectations, annotating cell populations using tissue-appropriate marker knowledge, and revising any of these choices when intermediate results demand it. Existing workflow systems such as Nextflow (Di Tommaso et al., 2017) and Snakemake (Köster and Rahmann, 2012; Mölder et al., 2021) can encode validation branches, QC checkpoints, and conditional logic. What they do not natively provide is open-ended adaptive reasoning: the ability to respond to analytical conditions they were not explicitly programmed to handle, propose a new integration strategy when the expected correction proves insufficient, or revise earlier decisions based on the qualitative character of downstream outputs. Manual expert analysis provides this reasoning but does not scale: one analyst cannot concurrently perform deep, iterative analyses of hundreds of datasets.

Large language models (LLMs) have introduced a new kind of tool into this space. Beyond code generation, modern LLMs embed substantial scientific knowledge and can reason through multi-step analytical problems. Foundational work on chain-of-thought prompting (Wei et al., 2022) and the ReAct observe–reason–act loop (Yao et al., 2023) demonstrated that LLMs reason more reliably when analytical steps are interleaved with execution rather than planned in a single pass. Multi-agent architectures such as AutoGen (Wu et al., 2023) and MetaGPT (Hong et al., 2023) extended this to structured division of cognitive labor among specialized agents. Domain deployments in chemistry, autonomous synthesis planning (Boiko et al., 2023) and tool-augmented molecular reasoning (M Bran et al., 2024), have shown that agentic LLMs can perform meaningful scientific work when given appropriate grounding. The concept of the AI scientist has moved rapidly from theoretical proposal to active engineering efforts.

We are past the point of asking whether LLMs can write bioinformatics code. The more pressing question is whether we can deploy them in actual research environments in ways that make them genuinely useful. This distinction brings up three requirements that laboratory benchmarks routinely omit. The first is *capability*: not raw model competence, but capability as implemented through a chassis that encodes domain-specific knowledge, grounds every analytical claim in real execution, and validates outcomes against explicit criteria. The model alone is not enough; the harness determines how much of the model’s capacity is directed toward useful analytical work. The second is *scalability*: the ability to run in actual institutional HPC environments without requiring workarounds. Production clusters prohibit Docker containers, a widely used software containerization platform, and outbound network connections by policy; any framework that requires these cannot be deployed where the data live. The third is *ingenuity*: the preservation of adaptive reasoning within a structured chassis. A system rigidly constrained to execute predetermined steps provides automation without intelligence. A system with no structural constraints produces unpredictable outputs that cannot be trusted for scientific conclusions. The engineering challenge is to constrain behavior enough to produce reliable, auditable analysis while leaving enough freedom for the system to detect anomalies, diagnose problems, and construct non-template solutions.

These three requirements are not independent. Over-constraining for capability and scalability risks eliminating ingenuity. Under-constraining preserves ingenuity but yields inconsistent outputs. Real research environments add further specificity: a research group’s analytical standards encode not just which tools to use but how to use them rigorously—the “taste” of the lab—which must be encoded into any AI assistant that is to be genuinely useful rather than merely technically compliant.

CARIBOU differs from adjacent systems in three ways that directly reflect these requirements. Unlike general multi-agent frameworks and LLM code assistants, it is HPC-native rather than notebook- or cloud-native, operating entirely through a Singularity-based backend that requires no network connections and is compatible with institutional cluster policies. Unlike workflow engines that encode fixed analytical decisions, CARIBOU represents agent behavior as researcher-modifiable blueprints, allowing lab-specific judgment and domain standards to be encoded and updated without model fine-tuning or pipeline rewriting. Also, unlike systems that invoke analytical tools statelessly, CARIBOU maintains a single persistent biological analysis state, represented as a living AnnData object, across interactions between multiple specialized AI agents responsible for different analytical tasks. These distinctions are summarized in the comparison presented in the Results.

To explore what a framework meeting these three requirements looks like in practice, we developed CARIBOU (Computational AI Research Interface for Bioinformatics, Omics, and Unifying Agents). CARIBOU is built around three core design principles. First, it uses researcher-editable agent blueprints that define analytical roles, domain-specific guidance, and patterns of collaboration between specialized AI agents without rigidly prescribing every step of analysis. Second, all analyses are executed within persistent and reproducible computational environments compatible with institutional HPC systems, including Singularity/Apptainer-based infrastructure commonly used in academic research. Third, CARIBOU continuously evaluates and refines its own analysis steps using feedback from the active computational workflow, allowing adaptive troubleshooting while restricting analyses to executable and validated operations. We present CARIBOU as an initial exploration of this design space, characterize its behavior across unit-task benchmarks, a metadata reconstruction challenge, and two end-to-end single-cell RNA-seq analyses, and offer both the architecture and its evaluation as contributions to the ongoing development of AI systems for computational biology.

## RESULTS

### The CARIBOU Framework: Architecture of an In-Silico Research Group

CARIBOU is implemented in Python 3.10 and operated through a command-line interface (CLI). The framework is organized into four integrated components. First, a researcher-editable agent configuration system defines the roles, responsibilities, and communication patterns of specialized AI agents assigned to different stages of analysis. Second, all generated code is executed inside an isolated and reproducible computational environment, ensuring that analyses are grounded in real execution rather than text-only responses. Third, CARIBOU maintains a persistent analytical context across long workflows, allowing intermediate results, dataset state, and prior decisions to remain available throughout multi-step analyses. Finally, the framework includes an automated benchmarking system for evaluating analytical performance and workflow behavior across standardized tasks (Figure 1).

**Figure 1.**
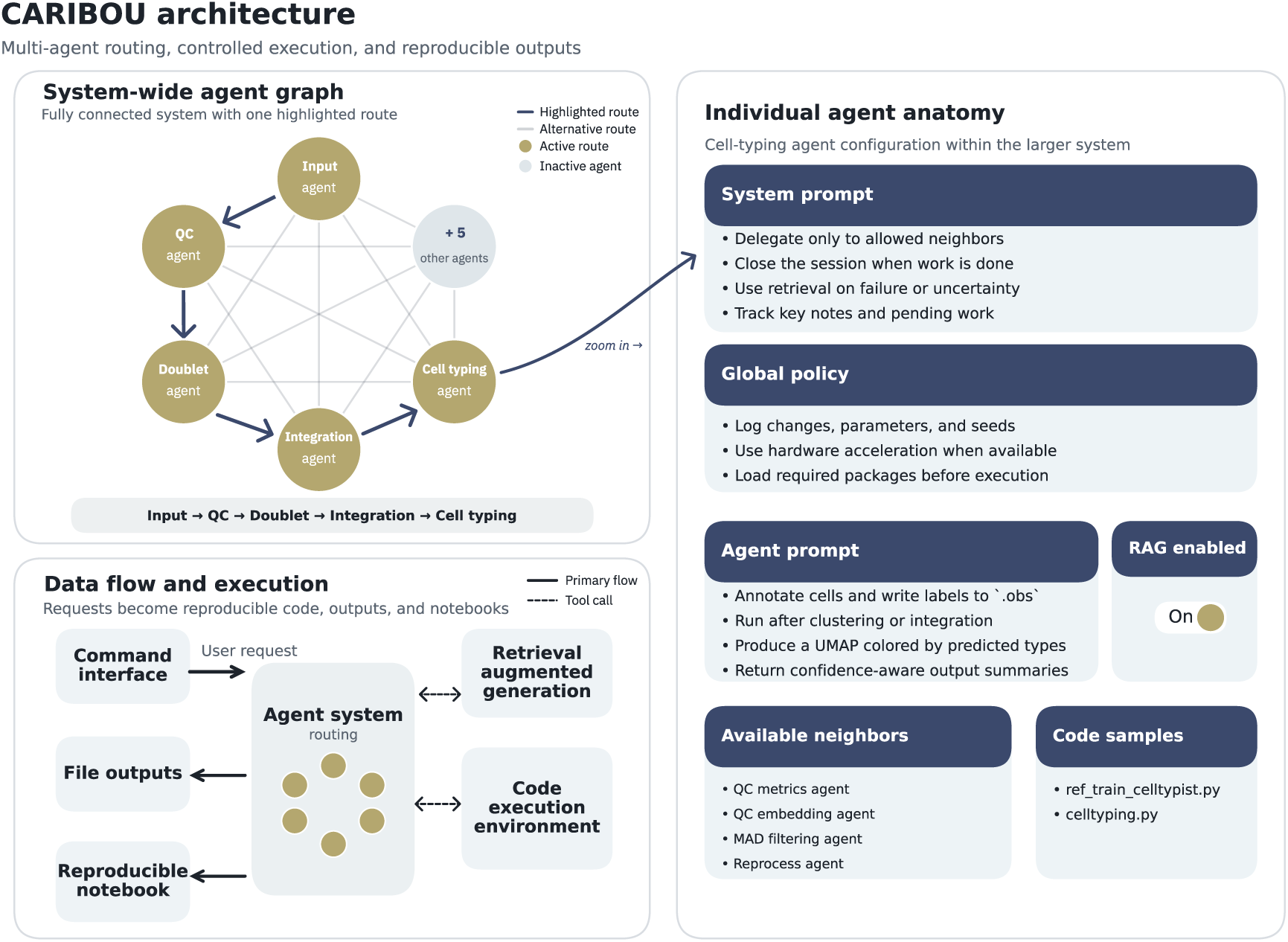
CARIBOU System Architecture. (*Top Left*) System-wide agent graph illustrating the fully connected multi-agent topology with one highlighted route from Input Agent through QC, Doublet, Integration, and Cell Typing agents. (*Center right*) Individual agent anatomy: system prompt with behavioral rules, global policy for shared analytical standards, agent-specific prompt, RAG enable/disable toggle, available neighboring agents, and referenced code samples. (*Bottom left*) Data flow and execution: user requests pass through the CARIBOU command interface to the agent system, which coordinates with a code execution environment and retrieval-augmented generation module, producing file outputs and a reproducible Jupyter notebook.

### Researcher-Configurable Multi-Agent Workflows

For single-cell RNA-seq analyses, CARIBOU uses a researcher-editable configuration blueprint that defines a virtual “in-silico research group” composed of multiple specialized AI agents working together on different stages of analysis (Figure 1). The primary blueprint used in this study, caribou_fully_connected_v2, includes a master orchestration agent alongside nine analysis specialists responsible for tasks such as data loading and validation, quality-control (QC) metric computation, median absolute deviation (MAD)-based outlier filtering, doublet detection (Wolock et al., 2019), QC embedding visualization, data reprocessing, downstream dimensionality reduction and clustering (McInnes et al., 2018; Traag et al., 2019), batch integration (Korsunsky et al., 2019; Lopez et al., 2018), and cell type annotation (Domínguez Conde et al., 2022). The fully connected communication structure allows any agent to consult or delegate work to any other agent, enabling the system to revisit earlier analytical stages when downstream results suggest that upstream processing decisions may need revision.

These blueprints are stored as human-readable JSON configuration files that define the analytical structure of the system without requiring modification of the underlying CARIBOU source code. Rather than writing procedural software from scratch for each experiment, researchers can modify templates that specify which agents are present, the analytical responsibilities assigned to each agent, how agents communicate with one another, and what domain-specific guidance or reference materials they can access. In practice, this allows CARIBOU to be adapted to different biological workflows by editing configuration files rather than rebuilding the framework itself (Supplemental Figure S1).

Each agent contains a role-specific instruction set describing its area of expertise and analytical responsibilities. Agents may also access curated code examples and function usage patterns for common bioinformatics tasks. These examples are provided as reference material rather than executable imported software modules, meaning that agents must generate and adapt their own code for the dataset under analysis instead of simply replaying fixed scripts. This design encourages analyses that remain grounded in established bioinformatics practices while preserving the flexibility needed to adapt workflows to different datasets, experimental designs, and biological questions.

During analysis, CARIBOU can switch between specialized agents while preserving the same active computational session and in-memory AnnData object. This allows intermediate results, dataset modifications, and analytical context to persist across specialist handoffs.

### Building for Scalability and HPC-Native Execution

CARIBOU executes all generated code inside an isolated and reproducible computational environment, allowing analyses to run consistently across different institutional systems. For the analyses in this paper, CARIBOU used a Singularity/Apptainer-based backend designed for high-performance computing (HPC) clusters where Docker, internet access, or networked services are often restricted (Kurtzer et al., 2017). This design allows CARIBOU to operate in the same security-constrained environments where large-scale biological datasets are commonly stored and analyzed (Supplemental Figure S4).

A key feature of this execution environment is that the analysis session persists across agent interactions. Rather than restarting the computational workspace each time an agent writes code or hands work to another specialist, CARIBOU maintains the same active Python session throughout the workflow. As a result, the same in-memory AnnData object persists across quality control, doublet detection, integration, clustering, and cell-type annotation. This creates a single evolving analysis state, closer to how an expert bioinformatician works interactively, rather than a sequence of disconnected tool calls.

The containerized environment includes a standardized bioinformatics software stack: Scanpy for single-cell analysis (Wolf et al., 2018), Scrublet for doublet detection (Wolock et al., 2019), HarmonyPy and scVI-tools for batch integration (Korsunsky et al., 2019; Lopez et al., 2018), and CellTypist for cell-type annotation (Domínguez Conde et al., 2022). By fixing this computational environment, CARIBOU improves reproducibility across runs and host systems while remaining compatible with institutional HPC infrastructure.

### Iterative Execution and Error-Corrective Workflow Behavior

CARIBOU is designed to prevent a common failure mode of LLM-based analysis systems: describing an analysis without actually performing it. Rather than allowing agents to respond only with narrative plans, CARIBOU requires each step to produce an actionable operation, such as executing code, delegating to another specialist, querying reference material, or ending the session. This keeps the system grounded in the active computational workflow.

Within this structure, agents still retain flexibility in how they respond to the data. They can generate new code, inspect execution output, revise failed analyses, delegate to another specialist, or choose a different analytical strategy when intermediate results suggest that the original path is insufficient. Thus, CARIBOU’s execution loop is constrained enough to require empirical testing of analytical claims, but open enough to preserve adaptive troubleshooting and non-template responses to unexpected data conditions.

This design supports an execute–observe–correct pattern: agents propose an analysis step, run it in the live computational environment, observe the result or error, and then decide how to proceed. As shown in the benchmarks and case studies below, this structure enables CARIBOU to recover from missing fields, failed function calls, and dataset-specific analysis requirements while maintaining a reproducible record of each step.

### Transparency Artifacts

Every CARIBOU run generates a structured set of transparency artifacts. A digital lab notebook is produced as a Jupyter notebook (Kluyver et al., 2016) containing the full agent conversation, all executed code cells, and all plot outputs, providing a complete, reproducible record of every decision made during analysis. A JSON session report captures runtime statistics. A structured TODO and NOTE system allows agents to emit persistent task lists and scientific observations that are tracked across agent transitions and saved to disk. For automated performance evaluation, CARIBOU exposes a pluggable benchmarking module interface, allowing researchers to define domain-specific quality metrics without modifying the core framework.

### Positioning CARIBOU Among Adjacent Systems

**Table 1.**
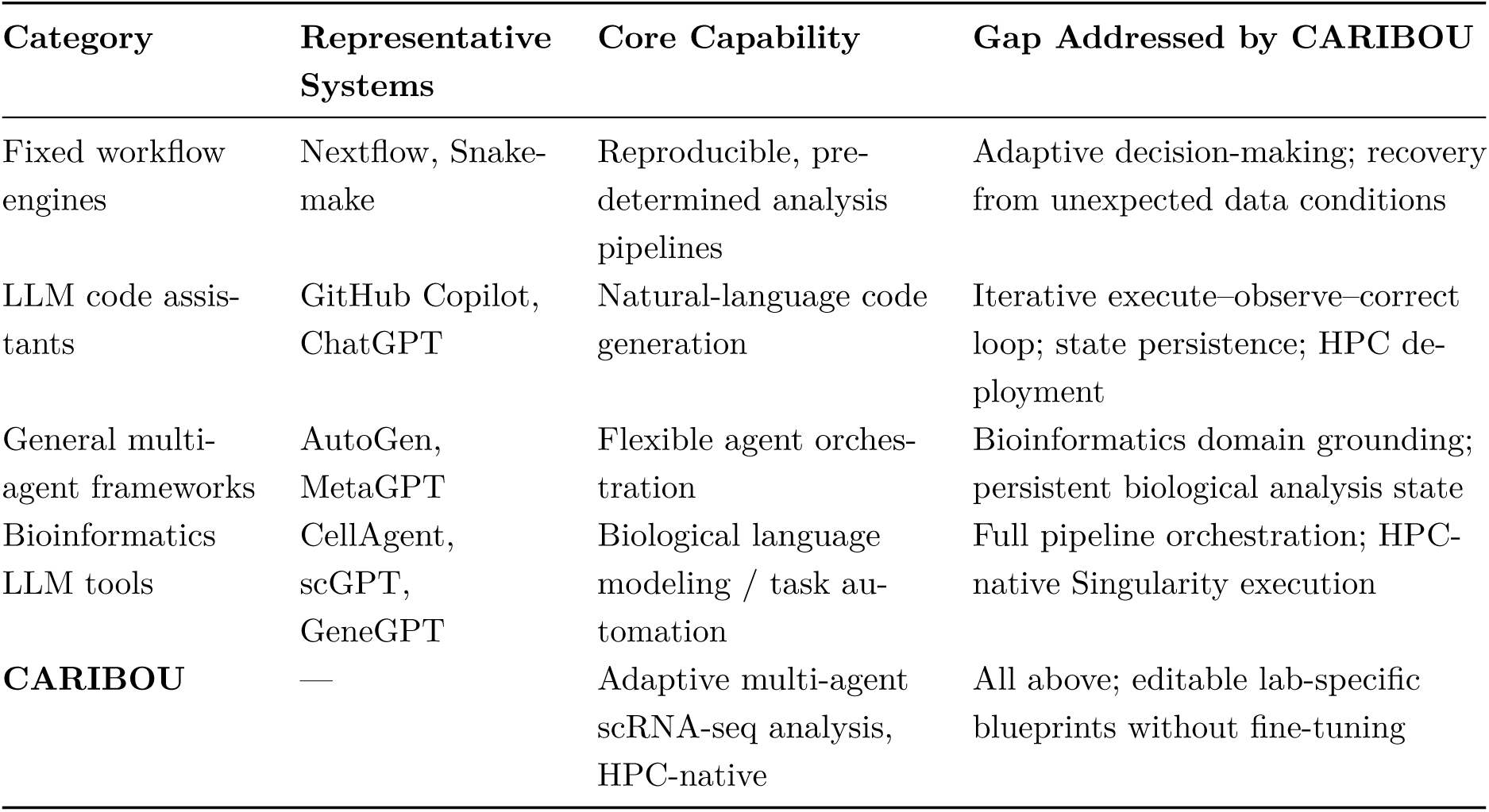
Comparison of CARIBOU with adjacent system categories. Existing categories each address one dimension of the problem—automation, code generation, or agent orchestration. CARIBOU is designed to combine HPC-native deployment, persistent biological analysis state, and researcher-tunable analytical judgment.

### Benchmarking Capability: Unit Task Performance

To characterize CARIBOU’s analytical capability on bounded tasks, we constructed a suite of unit benchmarks covering common scRNA-seq analytical steps: data loading and validation, full QC pipeline execution, doublet detection, and differential expression analysis (Figure 2A). Each task was evaluated across three execution configurations and two LLM backends (GPT-4o (OpenAI, 2023) and DeepSeek-Chat (DeepSeek-AI, 2024)).

**Figure 2.**
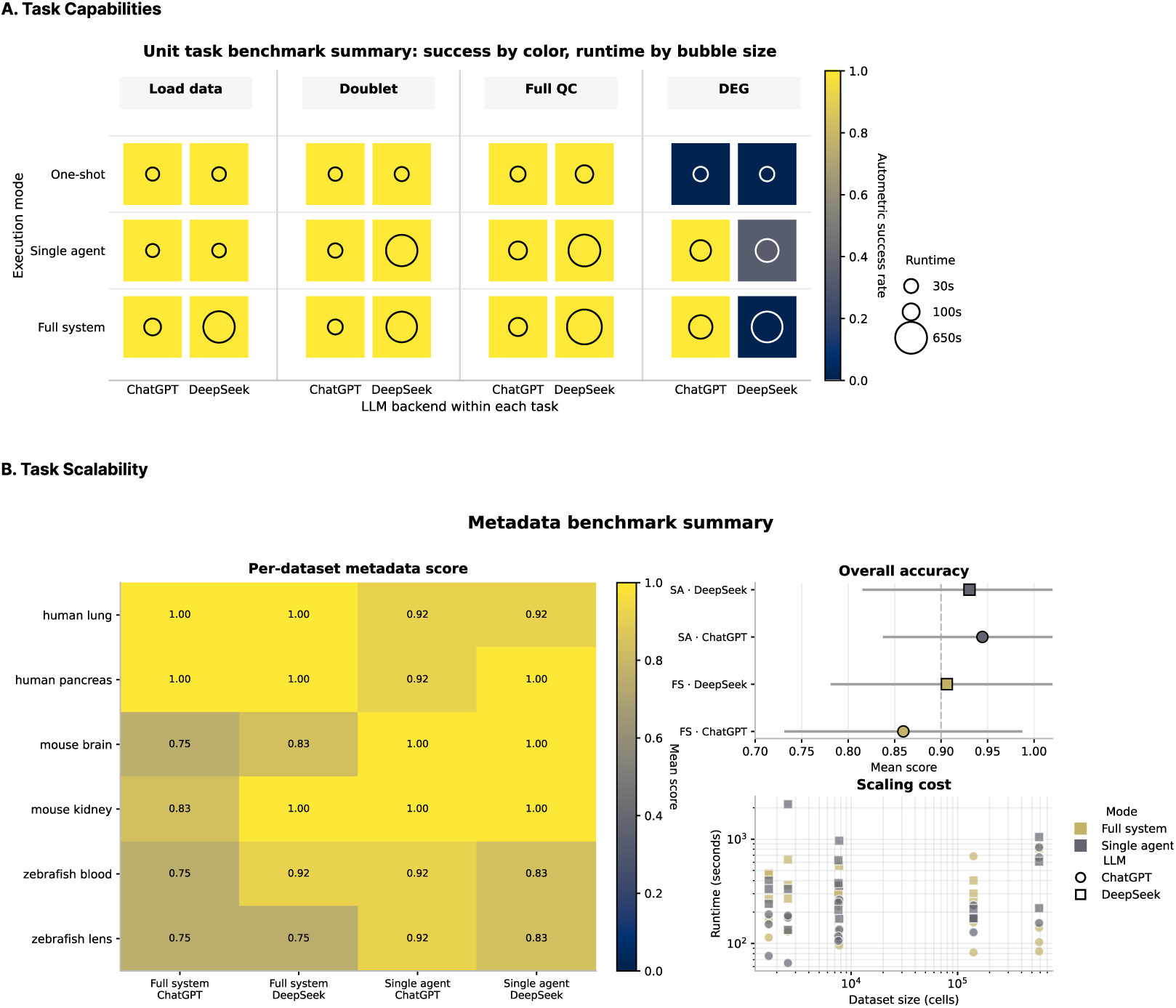
Unit Task and Scalability Benchmarking. (*A. Task Capabilities*) Unit task benchmark summary showing pipeline success (color: yellow = success, blue = failure) and total runtime (bubble size, seconds) across four task categories—Load Data, Doublet Detection, Full QC, and Differential Expression (DEG)—for three execution modes (One Shot, Single Agent, Full System) and two LLM backends (ChatGPT, DeepSeek). (*B. Metadata Reconstruction Benchmark*) Performance on metadata reconstruction across six anonymized public datasets (human lung, human pancreas, mouse brain, mouse kidney, zebrafish blood, zebrafish lens). Composite scores combine four metadata sub-tasks: species inference, organ inference, cell count estimation, and transcript count estimation. Species, cell count, and transcript metrics were near-perfect across all configurations; most performance variability arose from organ inference. Heatmaps show per-dataset composite accuracy, while accompanying plots summarize organ inference performance and runtime scaling with dataset size (cells, log scale).

The three configurations span increasing levels of structural support. One-shot mode submits a single LLM call without an iterative execution loop, analogous to querying an LLM for a code snippet without running or correcting it. Single-agent mode uses the iterative execution loop with a single specialist, allowing the agent to observe execution output, correct errors, and revise its approach. Full-system mode engages the complete multi-agent blueprint, enabling specialist delegation and the full collaborative workflow.

Success is assessed against explicit operational criteria rather than subjective judgment. The QC benchmark requires designated obs fields, a normalized counts layer, PCA with at least ten components, UMAP coordinates, and highly variable gene annotation. The batch correction benchmark requires measurable improvement in SCIB (Luecken et al., 2022) batch-mixing metrics without unacceptable degradation of cell-type structure. The differential expression benchmark validates cluster coverage, significant gene fraction, median absolute log fold-change, and top-gene specificity. These criteria are implemented as code-level checks within the benchmark harness, making success a matter of structural and quantitative contract, not interpretation.

Across task types and both model backends, the iterative execution loop outperforms one-shot completion. For complex multi-step tasks—full QC pipeline execution and differential expression— the full-system configuration achieves the most consistent pass rates. For bounded tasks such as data loading, single-agent performance matches full-system, suggesting that multi-agent delegation overhead is most justified where the workflow requires sustained, sequential decision-making across multiple analytical stages. Reported runtimes include LLM API latency, subject to provider-side variability and not comparable across providers.

An important caveat is that the current benchmark design does not cleanly isolate the contribution of multi-agent delegation from other architectural choices. All three conditions use the same underlying LLM and the same code-sample grounding; the single-agent condition retains the iterative execution loop and differs only in the absence of specialist delegation and the fully connected agent graph. The primary result—that the execution loop substantially outperforms one-shot completion—is robust. The secondary result—that full multi-agent delegation adds value over a single-agent loop—is consistent with the data but would benefit from a more controlled ablation. Table 2 summarizes which components are present in each tested condition.

**Table 2.** Architectural ablation of CARIBOU benchmark conditions.

| Condition | Persistent REPL | Execution loop | RAG | Multi-agent delegation | Blueprint code samples |
| --- | --- | --- | --- | --- | --- |
| One-shot | × | × | × | × | × |
| Single-agent loop | ✓ | ✓ | ✓ | × | ✓ |
| Full agent system | ✓ | ✓ | ✓ | ✓ | ✓ |

### Benchmarking Scalability: Metadata Reconstruction

Scalability in CARIBOU is addressed at the execution-substrate level; the HPC-native backend enables parallelization across datasets without Docker or network requirements. To provide a quantitative characterization of scalability across heterogeneous datasets, and to address a practically motivated use case, we constructed a metadata reconstruction benchmark (Figure 2B).

Many research institutions maintain archives of historical sequencing data that have been partially or fully stripped of provenance information through database migration, anonymization, or poor data management, leaving count matrices whose biological context is unclear. To evaluate whether CARIBOU could recover this missing context, we assembled a curated set of six public datasets and recorded ground-truth metadata fields including species, organ, cell count, and mean transcript count. We then generated anonymized “blind” versions of each dataset by removing stored annotations and analysis metadata from the AnnData object—including cell-level metadata tables, unstructured analysis notes, dimensionality reduction embeddings, graph relationships, and gene-level annotations— while preserving the underlying gene identifiers and count matrix. CARIBOU was then tasked with inferring the removed biological metadata using only the remaining expression data.

The metadata blueprint uses a staged serial workflow: an input agent characterizes the matrix structure, a species agent infers taxonomy from Ensembl identifier prefixes, an organ agent performs marker-based organ scoring using a deterministic 20,000-cell subset for large datasets to control runtime, and a summary agent writes the final inference. Scoring uses exact match for categorical fields and 5% relative tolerance for quantitative fields, providing objective success criteria that do not depend on interpretation.

CARIBOU achieved perfect or near-perfect performance on deterministic metadata recovery tasks, including species inference, cell count estimation, and transcript count estimation across all tested datasets. Variability in the composite metadata reconstruction score arose almost entirely from organ inference, which represents the biologically nontrivial component of the benchmark.

Organ classification performed reliably for tissues with highly distinctive marker profiles, such as human lung and pancreas, but degraded substantially for datasets with weaker or more ambiguous tissue-specific expression structure, including zebrafish lens and blood datasets. Across configurations, single-agent systems slightly outperformed the fully connected multi-agent system on this benchmark, suggesting that additional delegation overhead does not necessarily improve performance for bounded inference tasks with relatively narrow analytical scope.

These results indicate that CARIBOU can reliably recover structural and quantitative metadata from partially anonymized single-cell datasets while highlighting the current limits of marker-based biological inference in ambiguous tissue contexts. Runtime scaled with dataset size in a manner consistent with the computational cost of marker-based scoring and subset generation. Broader validation across additional tissues, disease states, and degraded datasets will be required to determine the generality of these findings.

### Natural Language to Reproducible Output: Prompt-to-Result Demonstrations

To illustrate CARIBOU’s adaptive analytical capabilities, we present three prompt-to-result case studies demonstrating end-to-end autonomous execution from natural language requests to publication-quality outputs (Figures 3–5). In Case 1, CARIBOU receives a request to generate a quality-control (QC) comparison report and autonomously loads pre- and post-QC AnnData objects, computes missing QC metrics, applies median absolute deviation (MAD)-based filtering thresholds, generates comparative visualizations, and assembles a final PDF report. The resulting figures summarize distributions for metrics including total transcript counts, detected gene counts, mitochondrial transcript percentage, and top-20 gene abundance before and after filtering.

**Figure 3.**
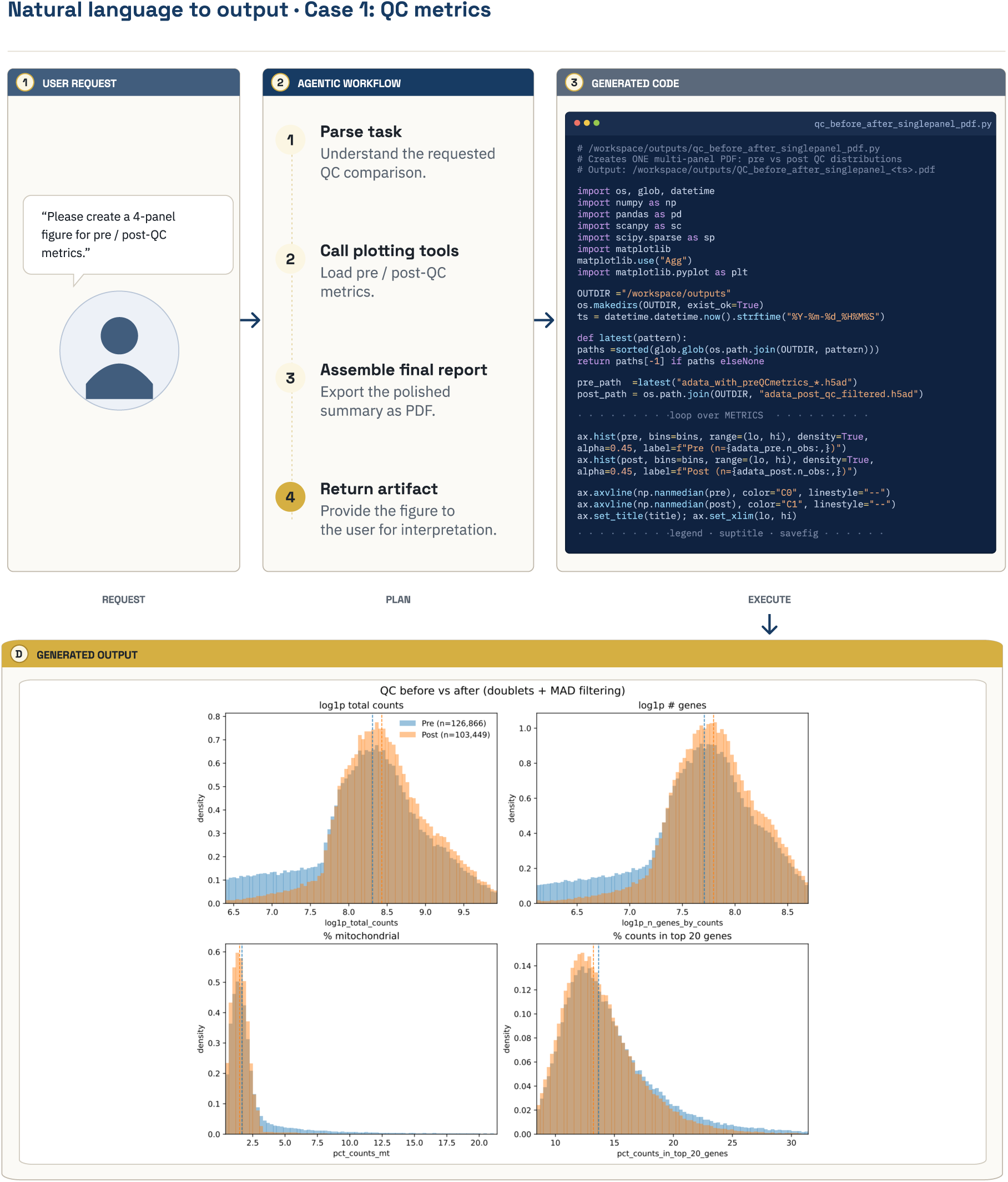
Prompt-to-Result Case Study 1: QC Metric Report. User requests QC metrics to be calculated pre/post QC with a figure package at the end of the task. CARIBOU parses the task and begins to call the necessary tools and agents. In this case the Input Agent, QC Agent, and MAD Agent are all necessary to load the data, identify the current metrics, calculate missing metrics, and then apply MAD for filtering. Code is then computed with an assembly of a figure package showing MAD filters applied before/after QC for user review.

**Figure 4.**
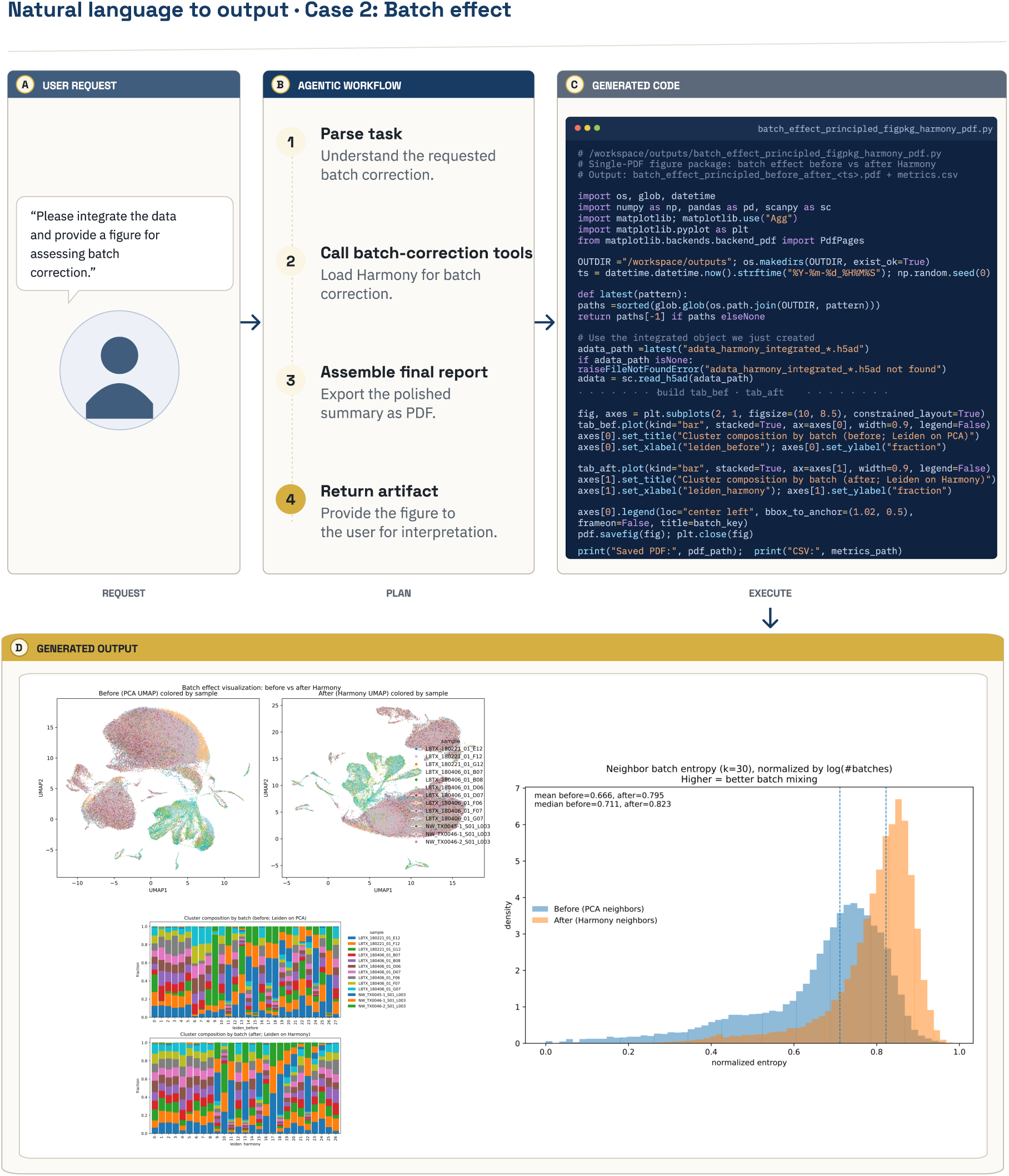
Prompt-to-Result Case Study 2: Batch Correction. User requests batch correction with a figure package at the end of the task. CARIBOU parses the task and begins to call the necessary tools and agents—in this case the Input Agent and Integration Agent. Harmony is called for batch correction and is applied to the data. A figure package is generated showing before/after Harmony integration, neighbor batch entropy as a mixing metric before/after Harmony integration, and finally Leiden mixing between batches before/after Harmony.

**Figure 5.**
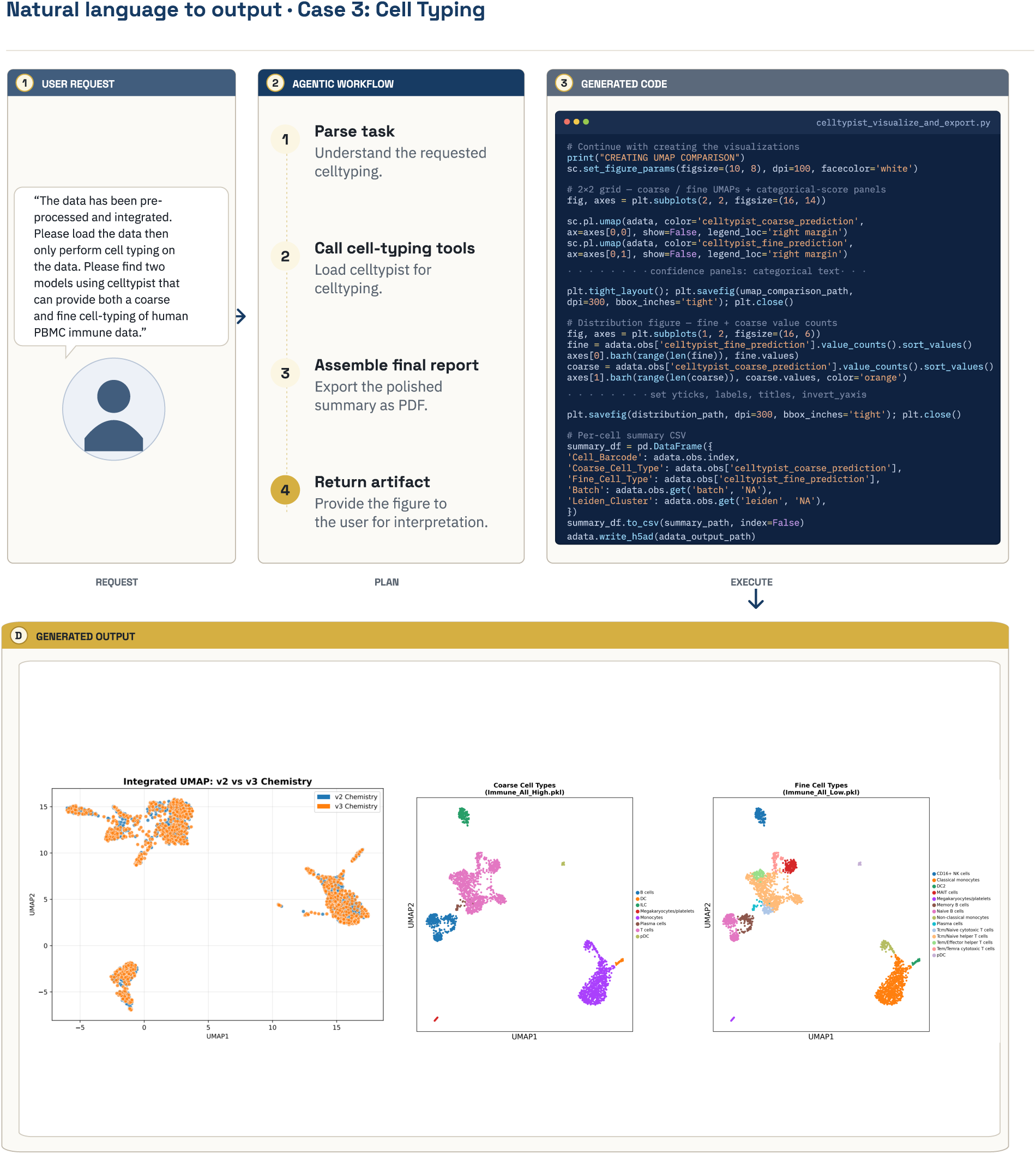
Prompt-to-Result Case Study 3: Cell-Type Annotation. User requests coarse and fine cell-type annotation for preprocessed and integrated PBMC data. CARIBOU parses the request, loads the integrated AnnData object, selects appropriate CellTypist immune reference models, performs multi-resolution cell typing, generates UMAP visualizations for batch chemistry, coarse cell types, and fine cell types, and exports both the figure package and per-cell annotation table for downstream review.

In Case 2, CARIBOU is tasked with assessing and correcting batch effects. Starting from the raw dataset, the system performs QC, identifies batch structure, selects an integration strategy appropriate to the observed data characteristics, executes Harmony or scVI-based integration, generates Uniform Manifold Approximation and Projection (UMAP) embeddings, and exports quantitative batch-mixing visualizations and summary figures. This case demonstrates CARIBOU’s ability to make sequential analytical decisions across multiple stages of a workflow rather than executing a fixed predefined pipeline.

In Case 3, CARIBOU performs automated cell-type annotation on Peripheral Blood Mononuclear Cell (PBMC) data using CellTypist-based annotation models. The system identifies appropriate annotation models for both coarse and fine-grained labeling schemes, performs annotation, generates visualization outputs, and exports labeled datasets for downstream analysis. This demonstrates the ability of CARIBOU to perform multi-resolution cell typing workflows commonly used in atlas-scale analyses.

Across all three cases, CARIBOU autonomously selects analytical tools, generates and executes code, iteratively corrects errors through the execute–observe–correct loop, and produces reproducible analysis artifacts without human intervention.

### End-to-End Analysis: Batch Integration Quality in the Allen Brain Atlas Hippocampus

The following two end-to-end analyses are intended as operational validation: they ask whether CARIBOU can execute a complete, biologically reasonable scRNA-seq pipeline from raw data to annotated output, not whether it makes novel biological discoveries. We make no new biological claims about the datasets used; the reference comparisons are measures of analytical reproducibility relative to manually curated processing of the same data.

To assess end-to-end analytical quality in an integration-heavy context, we applied CARIBOU in auto mode—a single natural language prompt followed by no subsequent human intervention—to the Allen Brain Atlas hippocampus dataset (Yao et al., 2023) (ABA-HIP; 85,955 cells). Starting from raw AnnData objects generated immediately after Cell Ranger (Zheng et al., 2017), the standard 10x Genomics preprocessing pipeline that converts raw sequencing reads into cell-by-gene count matrices, CARIBOU executed the complete pipeline through quality control, doublet detection, normalization, dimensionality reduction, batch integration, and cluster characterization. The full-system condition uses caribou_fully_connected_v2.json; the single-agent condition uses caribou_single_agent.json.

Integration quality was assessed using SCIB metrics, a standardized framework for evaluating single-cell data integration quality (Luecken et al., 2022), including average silhouette width (batch), graph connectivity, and integration local inverse Simpson’s index (iLISI). An important methodological note: bio-conservation metrics use reference cell-type labels projected onto aligned CARIBOU outputs rather than CARIBOU’s own predicted labels. This makes the integration benchmark a measure of structural preservation independent of labeling style—the question is whether CARIBOU produces a latent space that respects the underlying biology, not whether it uses the same vocabulary as the reference. Reference baselines were established from manually curated processing of the same data.

Across GPT-4o and DeepSeek-Chat backends, CARIBOU’s full-system outputs achieved integration metrics within the range produced by the reference pipeline, with batch mixing and cell-type structure preservation comparable to the curated reference. The single-agent condition showed greater variability, consistent with the hypothesis that sequential specialist delegation improves analytical coherence across the cascading decisions of a multi-step integration workflow. Neither model backend consistently dominated across all metrics, and run-to-run variation reflects the stochastic nature of LLM-generated analytical decisions. These results are best interpreted as a feasibility demonstration on a single benchmark dataset; broader characterization across diverse biological contexts is required.

### End-to-End Analysis: Cell Type Annotation in Tabula Sapiens Large Intestine

To assess end-to-end analytical quality in a cell-typing context, we applied CARIBOU in auto mode to the Tabula Sapiens large intestine dataset (Tabula Sapiens Consortium et al., 2022), comparing predicted cell-type annotations against authoritative consortium labels across 12 major cell populations. Starting from raw reads processed to AnnData, CARIBOU executed the full pipeline through QC, normalization, clustering, and CellTypist-based annotation (Domínguez Conde et al., 2022). The same blueprint pair as the ABA-HIP benchmark was used.

Cell-type annotation accuracy was assessed using several complementary metrics comparing CARIBOU-generated labels against reference annotations. Weighted F1 score measures the balance between precision and recall across cell types while accounting for differences in population size. Adjusted Rand index (ARI) evaluates how similarly cells are grouped into clusters relative to the reference annotation, correcting for agreement expected by chance. Normalized mutual information (NMI) measures the overall similarity between predicted and reference cluster assignments as an information-sharing problem, with higher values indicating stronger agreement between labeling structures. PCA k-nearest-neighbor overlap assesses whether cells remain close to biologically similar neighbors in reduced-dimensional expression space after analysis, providing a measure of local structural preservation within the dataset. Label harmonization was performed through dataset-specific mapping dictionaries to prevent vocabulary mismatches, differences in cell-type naming conventions rather than differences in biological assignments, from dominating the quantitative scores.

For the Tabula Sapiens large intestine benchmark, CARIBOU produced cell-type annotations broadly aligned with consortium reference labels across all tested system configurations. Weighted F1 scores were highest for the DeepSeek single-agent configuration (0.88), followed by both the ChatGPT single-agent and DeepSeek full-system configurations (0.86), and finally the ChatGPT full-system configuration (0.84). Across configurations, ARI and NMI values remained consistently high, indicating that CARIBOU generally preserved the structure of major cell populations even when exact label assignments varied between runs or model backends.

Performance was strongest for abundant populations, such as colonocytes, T cells, and fibroblasts, and degraded for rare populations including tuft cells and mast cells. This pattern likely reflects the inherent difficulty of rare-cell annotation for reference-model-based approaches such as CellTypist rather than a limitation specific to CARIBOU itself. Overall, these results suggest that CARIBOU can reproduce expert-like annotations for well-characterized and abundant cell populations in familiar tissue contexts while highlighting remaining challenges associated with rare-cell classification and annotation consistency across heterogeneous biological datasets.

### Adaptive Reasoning: Non-Template Responses to Unexpected Data

The benchmarks above assess task completion against defined criteria. The most interesting property of an iterative execution system is what happens when the expected analytical path is unavailable, when the data do not conform to the template anticipated by the blueprint, or when execution failures require the workflow to dynamically adapt. Figure 8 presents a representative case study illustrating adaptive behavior within the CARIBOU execution loop.

The central case involves a natural language request to plot specific QC metrics. Upon executing the initial plotting code, the agent received an error indicating that the requested fields were absent from the AnnData object because they had not been computed during the upstream QC pass. Rather than failing, returning an error to the user, or silently plotting available metrics, the agent diagnosed the missing fields, generated the code required to compute them from the available count data, executed that code to populate the missing obs columns, and then proceeded with the original plotting request. The sequence detect–diagnose–generate–execute–continue was not explicitly hard-coded into the blueprint and instead emerged through iterative interaction between the language model and the execution environment.

We are careful not to overinterpret this behavior. These cases represent bounded instances of adaptive error recovery within structured analytical contexts rather than demonstrations of general autonomous scientific reasoning. Nevertheless, across both the primary example and the additional supplemental case studies, CARIBOU consistently demonstrates the ability to revise analytical plans in response to execution feedback rather than rigidly following predetermined workflows. This distinction between scripted execution and execution-conditioned adaptation becomes increasingly important as such systems are applied to heterogeneous real-world biological datasets.

## DISCUSSION

We have described CARIBOU as an initial exploration of the design space for LLM-based bioinformatics automation that works within the constraints of real institutional research environments. The three-property framework we introduced is not a claim about what CARIBOU has solved. It is a way of naming the design tradeoffs that any practical system must navigate. The architectural decisions we describe represent one position within that space.

The blueprint architecture encodes capability through researcher-modifiable specifications of role, knowledge, and analytical behavior rather than through fine-tuned model weights. This makes CARIBOU transparent and adaptable: researchers can inspect what guidance the system receives, modify its analytical expectations, and extend its capabilities without retraining the underlying language model. The limitation is that CARIBOU’s behavior remains bounded by the quality and coverage of its blueprints, code examples, and underlying model. Tasks not represented in the current blueprint—such as rigorous differential expression, gene ontology analysis, trajectory inference, or pathway analysis—may fail or require substantial human guidance.

This boundary is especially important for iterative analytical decisions. CARIBOU can execute and troubleshoot workflows, but it does not yet systematically optimize intermediate outputs such as QC thresholds, batch-corrected embeddings, or SCIB scores across repeated attempts. Expert bioinformaticians routinely refine these choices by interpreting artifacts such as ambient RNA contamination, doublet structure, over-filtering, and residual batch effects. Supporting this kind of iterative tuning at scale will require future versions of CARIBOU to incorporate explicit checkpointing, output validation, and automated refinement loops rather than relying primarily on conversational correction.

The long-lived REPL design addresses scalability and supports analytical continuity, but persistent state is also the framework’s most significant architectural liability. Early-pipeline errors propagate silently into downstream results, and the current framework has no systematic checkpoint-and- rollback mechanism to detect or recover from them. This is discussed further in the Failure Modes section below.

The benchmark results should be interpreted conservatively. The unit task benchmarks demonstrate consistent performance advantages for iterative execution compared to one-shot completion, which is a finding about the value of the execution loop itself, not a comparative claim about multi-agent architectures specifically. The two end-to-end benchmarks each use a single dataset and two model backends; performance across the full distribution of real-world scRNA-seq datasets—spanning diverse tissue types, organisms, sequencing protocols, and quality profiles—is unknown and is not addressed by the evidence presented here. The metadata reconstruction benchmark provides initial evidence for a practically motivated use case and demonstrates that the framework can be applied to novel task types beyond standard scRNA-seq analysis, but similarly requires wider validation.

The ingenuity case study illustrates a specific kind of adaptive behavior—error-driven field generation—that is enabled by the constrained execution loop. We present it as an illustration of what the design enables rather than as evidence of broad autonomous scientific reasoning. A rigorous characterization of adaptive behavior—when it occurs, when it fails, and what distinguishes the two—would require systematic evaluation that is beyond the scope of the present work.

### Failure Modes and Current Boundaries

We observed several recurring failure modes that define CARIBOU’s current practical boundaries.

#### Rare cell type annotation failures

CellTypist-based annotation systematically underperforms on rare populations (e.g., tuft cells, mast cells, enteroendocrine cells) where marker evidence is sparse. This is a limitation of the annotation method, not CARIBOU-specific, but it means CARIBOU inherits the accuracy ceiling of the tools it calls.

#### Organ ambiguity in metadata reconstruction

The metadata benchmark degrades for tissues with overlapping marker profiles—for example, distinguishing kidney cortex from kidney medulla, or distinguishing intestinal segments—where marker-panel-based scoring produces ambiguous results. The system does not reliably detect its own uncertainty in these cases and may return a confident but incorrect inference.

#### Cascading errors from early analytical decisions

The persistent analysis state that enables CARIBOU’s continuity is also its most significant architectural liability. An overly aggressive QC filtering step early in the session removes cells that cannot be recovered; an incorrect normalization choice propagates through every downstream analysis. The fully connected delegation graph allows an agent to revisit earlier stages, but this is an opportunistic escape hatch, not a systematic checkpoint-and-rollback mechanism. The framework currently has no way to automatically detect that a downstream anomaly is caused by an upstream decision rather than by a biological characteristic of the data. This is not a minor limitation: it means that CARIBOU sessions can produce internally consistent but analytically incorrect outputs without triggering any structural failure signal. Implementing explicit intermediate checkpoints, state snapshots, and rollback capabilities is a design priority, not merely a future improvement.

#### Stochastic run-to-run variability

Large language models generate responses through probabilistic sampling, meaning that the same analysis can produce slightly different decisions across independent runs. This variability can influence choices such as clustering resolution, filtering thresholds, or the ordering of quality-control steps, and is reflected in the spread of SCIB metrics observed across repeated analyses. As a result, CARIBOU does not guarantee fully deterministic outputs, and the downstream biological significance of this analytical variability remains to be systematically characterized.

#### Blueprint quality dependence

The system’s capability is bounded by the quality of the blueprint provided. A blueprint with incomplete agent coverage, poor code-sample grounding, or ambiguous role boundaries will produce degraded analytical outputs. There is no automated mechanism for detecting blueprint adequacy; this currently requires domain expert evaluation.

#### No guarantee of biological correctness

CARIBOU executes code against data and validates outputs against structural criteria, but it does not have access to biological ground truth. A session can complete successfully, passing all structural checks, while producing biologically incorrect conclusions. Expert review of CARIBOU outputs remains essential before drawing scientific conclusions.

#### Several directions warrant future investigation

A systematic comparison of context management strategies across session lengths and task types would clarify when different memory approaches provide meaningful advantages over the context limit compression strategy used in this work. In rolling window compression, older portions of the conversational history are periodically summarized while a recent uncompressed interaction window is retained to preserve short-term analytical continuity. In agent handoff compression, delegation events trigger the generation of a structured semantic summary by the outgoing agent, which is then passed to the incoming specialist in place of the full prior conversation history. Evaluating how these approaches influence analytical consistency, long-horizon reasoning, and error recovery remains an important direction for future work.

Extending CARIBOU with explicit output validation logic—including checks that generated AnnData objects satisfy expected biological and statistical properties—would further tighten the connection between execution and correctness beyond what is currently enforced through blueprint-level guidance alone. More broadly, evaluation across a wider range of biological domains, organisms, sequencing modalities, and analytical tasks will be necessary to establish the generality boundaries of the current framework.

CARIBOU’s session artifact system, including persisting run reports, benchmark ledgers, code snippets, notebook logs, and extracted notes, provides a technical foundation for auditability. However, the biological conclusions drawn from AI-generated analyses require expert evaluation, and community norms for attributing and validating AI-assisted scientific work are still being established. We offer CARIBOU as a contribution to that conversation, and we welcome scrutiny of both its architecture and its claims.

## METHODS

### CARIBOU Framework Architecture

CARIBOU is implemented in Python 3.10 as a command-line multi-agent execution framework. The user-facing interface is a CLI built with the Typer library, exposing commands for creating agent systems, running analyses, managing datasets, and configuring credentials. The framework’s central abstraction is a JSON blueprint that specifies: (i) a global analytical policy applied to all agents; (ii) a set of named agents, each with a role-specific system prompt; (iii) a directed delegation graph; (iv) optional retrieval-augmented grounding support; and (v) optional code-sample attachments for prompt grounding. Code samples are injected as reference text rather than imported as executable modules, requiring agents to generate and adapt analysis code dynamically for the dataset under study.

The primary blueprint for multi-agent scRNA-seq analysis is caribou_fully_connected_v2.json, defining nine specialists plus a master orchestrator. The single-agent condition uses caribou_ single_agent.json. For metadata benchmarks, the full-system condition uses dataset_metadata_ agent.json and the single-agent condition uses metadata_single_agent.json. Several bounded single-agent task runs use dedicated task-specific blueprints (qc_single_agent.json, load_single_ agent.json, deg_single_agent.json, etc.).

At runtime, the driver agent receives an initial context from two pinned system messages: one containing the global policy and one containing the current agent prompt plus the mounted dataset path. Delegation is mediated through regular-expression matching of command strings (e.g., delegate_to_Celltyping), replacing the active agent prompt without interrupting the session. The execution loop recognizes four action classes: executable Python code blocks, delegation commands, retrieval queries, and an explicit session terminator (end_session). Responses lacking a recognized action receive a corrective system injection instructing the model to execute, delegate, or retrieve rather than narrate analysis steps without action.

### Retrieval-Augmented Grounding

CARIBOU implements a lightweight retrieval-augmented generation (RAG) module designed primarily for function-signature recovery and code grounding during analysis execution. Retrieval support can be enabled on a per-agent basis within blueprint configuration files.

The retrieval system operates over a local JSON-based reference store containing curated code examples, function signatures, and usage patterns for supported bioinformatics tooling. When code execution fails with a recognizable function-signature or argument error, the orchestration layer automatically embeds the failed query using Qwen3-Embedding-0.6B (Zhang et al., 2025) and retrieves semantically similar entries from the reference store. Retrieved entries are injected back into the active agent context as supplemental reference material, allowing the agent to revise and regenerate code using corrected function usage patterns.

Agents may also issue explicit retrieval commands using the query_rag_<TOPIC> syntax recognized by the execution loop. Retrieved content is treated as advisory reference text rather than executable code and must still be rewritten and executed within the active sandbox environment. This preserves the execute–observe–correct structure of the framework while reducing failure rates caused by incorrect function usage or parameter specification.

The retrieval system is intentionally narrow in scope. It is not designed as a general biological knowledge base or internet-scale document retrieval system. Instead, it functions as a lightweight local grounding mechanism focused on improving execution reliability for bioinformatics workflows. If the embedding model is unavailable or retrieval initialization fails, CARIBOU degrades gracefully by disabling retrieval while preserving all other framework functionality.

### Execution Sandbox

All CARIBOU-generated code executes within an isolated and reproducible containerized execution environment built around Python 3.10 and a standardized bioinformatics software stack. The environment includes Scanpy (Wolf et al., 2018) for single-cell analysis, Scrublet for doublet detection (Wolock et al., 2019), HarmonyPy and scVI-tools for batch integration (Korsunsky et al., 2019; Lopez et al., 2018), and CellTypist for cell-type annotation (Domínguez Conde et al., 2022). By standardizing the execution environment, CARIBOU reduces variability across host systems and improves reproducibility of generated analyses.

CARIBOU supports two execution backends. The Docker backend runs a persistent Jupyter kernel exposed through a FastAPI interface, allowing generated code to execute while preserving Python state between calls. The Singularity/Apptainer backend (Kurtzer et al., 2017), used for all analyses in this paper, is designed for deployment on institutional high-performance computing (HPC) systems where Docker and outbound network communication are frequently restricted by policy.

In Singularity/Apptainer mode, CARIBOU launches a long-lived Python REPL inside the container, allowing the same Python interpreter and in-memory AnnData object to persist across all agent interactions during a session. Communication between the orchestration layer and execution environment occurs through structured stdin/stdout transaction boundaries. This persistent execution model enables analytical continuity across multiple stages of analysis—including quality control, doublet detection, integration, clustering, and cell-type annotation—without reinitializing the computational state between agent transitions.

Input datasets and output directories are exposed to the container through explicitly mounted filesystem paths, restricting generated code access to designated analysis locations. In HPC mode, analyses execute without outbound network communication, enabling deployment in security-restricted institutional computing environments where large-scale biological datasets are commonly stored. GPU acceleration is conditionally enabled when compatible NVIDIA CUDA devices are available, with blueprint agents instructed to preferentially use GPU-supported workflows when appropriate.

### Context Management

Three context management strategies are implemented. Context limit compression (used in all analyses in this paper) maintains the full growing transcript until the model token limit is approached, compresses all history into a single summary, and continues. Agent handoff compression generates a structured handoff report when an agent delegates, seeding the incoming agent’s context with a semantic summary; when enabled, episodic compression is disabled. Rolling window compression periodically compresses older message chunks while retaining a recent uncompressed tail. The full_system label in benchmark figures refers to the full multi-agent system with context limit compression.

### Unit Task Benchmarks

Unit tasks—data loading, full QC pipeline, doublet detection, and differential expression—were evaluated across one-shot, single-agent, and full-system configurations with GPT-4o and DeepSeek-Chat backends. One-shot runs submit a single LLM call without an execution loop. Single-agent runs use dedicated single-task blueprints for bounded tasks or caribou_fully_connected_v2.json with master_agent as driver for longer tasks. Full-system runs use caribou_fully_connected_v2.json.

Success criteria are task-specific operational contracts: the QC benchmark requires designated obs fields (total counts, mitochondrial fraction, log-normalized counts), a counts layer, PCA with *≥*10 components, UMAP coordinates, and highly variable gene annotation; batch correction requires measurable SCIB batch-mixing improvement without loss of cell-type structure; differential expression requires cluster coverage, significant gene fraction, median absolute log fold-change, and top-gene specificity. Reported runtimes include LLM API latency, subject to provider-side variability.

### Metadata Reconstruction Benchmark

Six public datasets were downloaded, ground-truth metadata fields were recorded, and anonymized versions were generated by removing obs, uns, obsm, obsp, and varm while preserving gene identifiers and count data. CARIBOU was challenged to infer species, organ, cell count, and mean transcript count. The staged metadata blueprint uses the following: input agent for matrix characterization, species agent for taxonomy inference from Ensembl identifier prefixes, organ agent for marker-panel-based scoring using a deterministic 20,000-cell subset for large datasets, and summary agent for final inference. Scoring uses exact match for categorical fields and *≤*5% relative tolerance for quantitative fields.

### End-to-End Benchmarks

For ABA-HIP integration benchmarks, 10x Chromium v2 and v3 hippocampus data were processed from raw AnnData (post-Cell Ranger) through the full CARIBOU pipeline in auto mode. Integration was assessed using SCIB metrics (Luecken et al., 2022): average silhouette width (batch and cell type), graph connectivity, and iLISI. Bio-conservation metrics use reference cell-type labels projected onto aligned CARIBOU outputs. Reference baselines were established by manually curated processing of the same data. Full-system runs use caribou_fully_connected_v2.json; single-agent runs use caribou_single_agent.json.

For Tabula Sapiens large intestine cell-typing benchmarks, raw reads were processed to AnnData then analyzed through the full CARIBOU pipeline in auto mode. Cell-type accuracy was assessed using weighted F1, ARI, NMI, and PCA k-NN overlap against Tabula Sapiens consortium labels. Label harmonization used dataset-specific mapping dictionaries. The same blueprint pair as ABA-HIP was used.

### Model Configurations

All benchmark experiments reported in this study were conducted using GPT-4o (OpenAI, 2023) (OpenAI) and DeepSeek-Chat (DeepSeek-AI, 2024) (DeepSeek) as the evaluated LLM backends. Integration, cell-typing, metadata reconstruction, and unit-task benchmarks used these provider-hosted models exclusively, and all figures and tables report the specific backend used for each run. One-shot benchmark experiments used GPT-4o via the OpenAI API.

CARIBOU additionally supports alternative inference backends, including Anthropic Claude models and locally hosted Ollama-compatible models. Local inference allows analyses to be executed without transmitting prompts or dataset-derived context to external API providers, which may be desirable for institutionally sensitive or network-restricted deployments. In these configurations, model calls are served through a locally hosted Ollama endpoint rather than a remote API service. These deployment pathways are supported by the framework but were not evaluated in the benchmarks presented here.

### Execution Constraints and Partial Isolation

Because CARIBOU autonomously executes generated code, the isolation properties of the execution environment are relevant to its deployment. We describe the current state accurately: the environment provides partial isolation enforced primarily through bind-mount scoping and blueprint-level policy, not through a formally verified security model.

#### File system access

In both Docker and Singularity backends, the executing container has access only to explicitly bind-mounted paths: the input dataset, the output directory, and optionally a CellTypist model cache. Files outside these mount points are not accessible to generated code. Agents are instructed in their blueprints to write outputs only to /workspace/outputs/.

#### Network access

In HPC (Singularity) mode, the container is launched without network interfaces, eliminating the possibility of generated code making outbound connections. In Docker mode, the container operates on a localhost-only bridge network; agents are instructed not to make external network calls, though this is enforced by policy rather than by a hard network rule in the current implementation.

#### Shell access and destructive operations

Agents generate Python code executed by a Python interpreter; they do not have direct shell access. Blueprint prompts include a policy-level instruction discouraging subprocess calls and file deletion outside the output directory. This is a soft constraint: Python code could in principle invoke os.system or subprocess, and future versions should implement a code-level allow-list to enforce these restrictions more rigorously.

#### Resource consumption

No hard memory or CPU limits are imposed by the framework itself; container-level resource limits (e.g., SLURM memory and time allocations) provide the primary constraint in HPC deployments. LLM API call budgets are bounded by the per-session turn limit and the context compression strategy. Infinite loops in generated code would eventually be terminated by the SLURM time limit or the LLM API timeout, but not by an internal CARIBOU watchdog in the current implementation.

#### Credential handling

API keys and provider credentials are read from environment variables or a local configuration file at CLI startup and are not injected into agent context, generated code, or session artifacts.

Local-model execution reduces dependence on external LLM APIs but does not change the code-execution isolation model. The generated Python still runs in the containerized backend, and the same file-system, subprocess, and resource-consumption boundaries apply.

### Reproducibility

All sessions automatically persist: a structured session report (turn count, execution attempts, execution failures, correction count, duration, end reason), a JSON benchmark ledger, a code snippets directory with all generated and executed code, a notebook-format chat log (Kluyver et al., 2016), and extracted NOTE and TODO artifacts under the run output directory. Together these artifacts allow post-hoc reconstruction of every analytical decision made during a session.

For the benchmark analyses in this paper, LLM sampling temperature was not explicitly set and defaults to the provider API default (temperature = 1 for GPT-4o; temperature = 1 for DeepSeek-Chat). As a result, individual runs are not deterministic and replicate runs will produce quantitatively similar but not identical outputs. The spread of SCIB metrics visible in Figure 6 reflects this inherent stochasticity. Exact model versions, blueprints, and benchmark launch scripts are released with the codebase on GitHub. All generated code from benchmark runs is archived in the per-session snippets/ directory.

**Figure 6.**
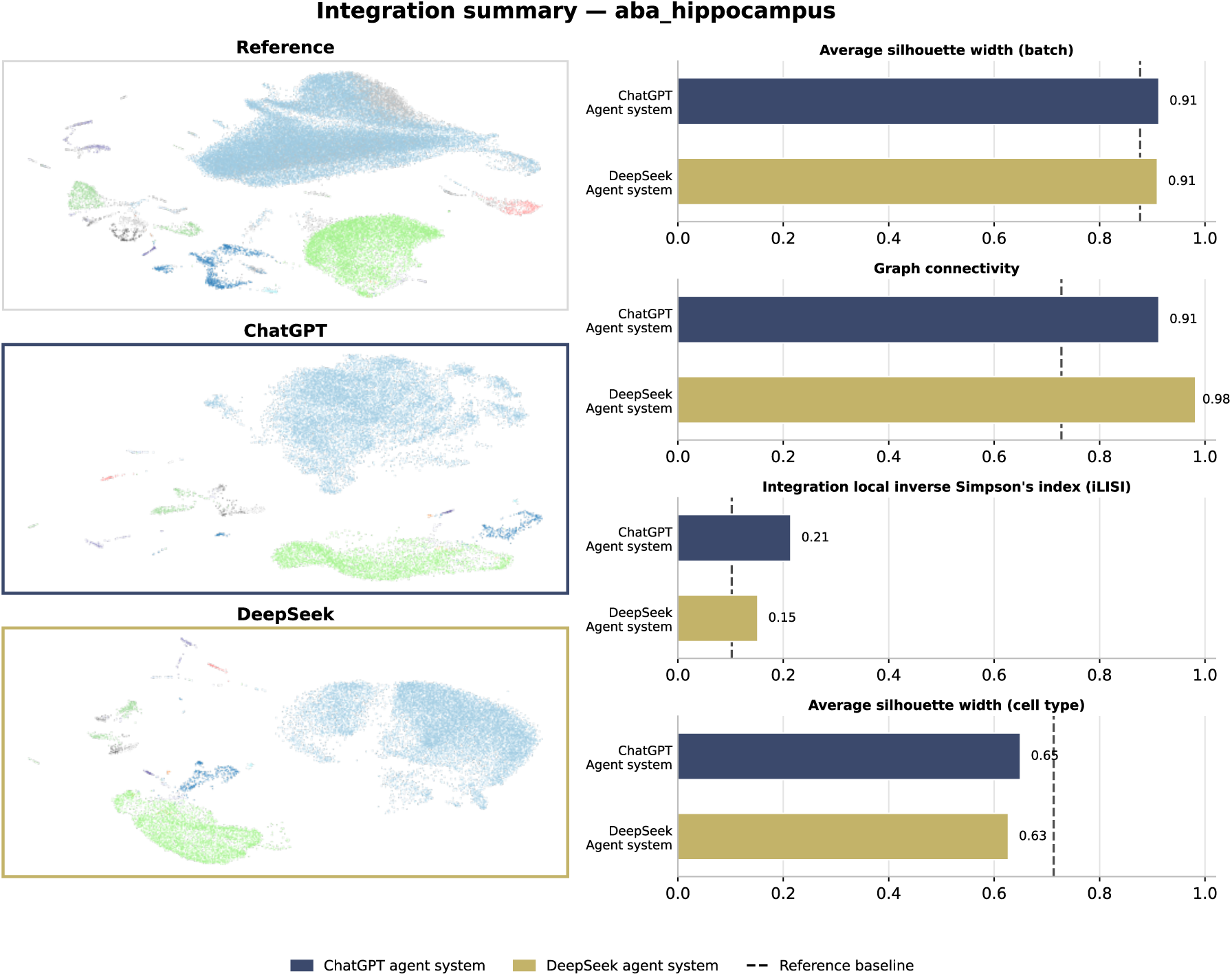
Integration Quality, Allen Brain Atlas Hippocampus. (*Left panels*) UMAP embeddings of the ABA-HIP dataset (85,955 cells, **13** batches) for the reference processing, the ChatGPT agent system, and the DeepSeek agent system, colored by major cell class. (*Right panels*) Quantitative SCIB integration metrics (Luecken et al., 2022) comparing ChatGPT and DeepSeek agent systems against the reference baseline: average silhouette width (batch), graph connectivity, integration local inverse Simpson’s index (iLISI), and average silhouette width (cell type). The dashed line indicates the reference baseline value.

**Figure 7.**
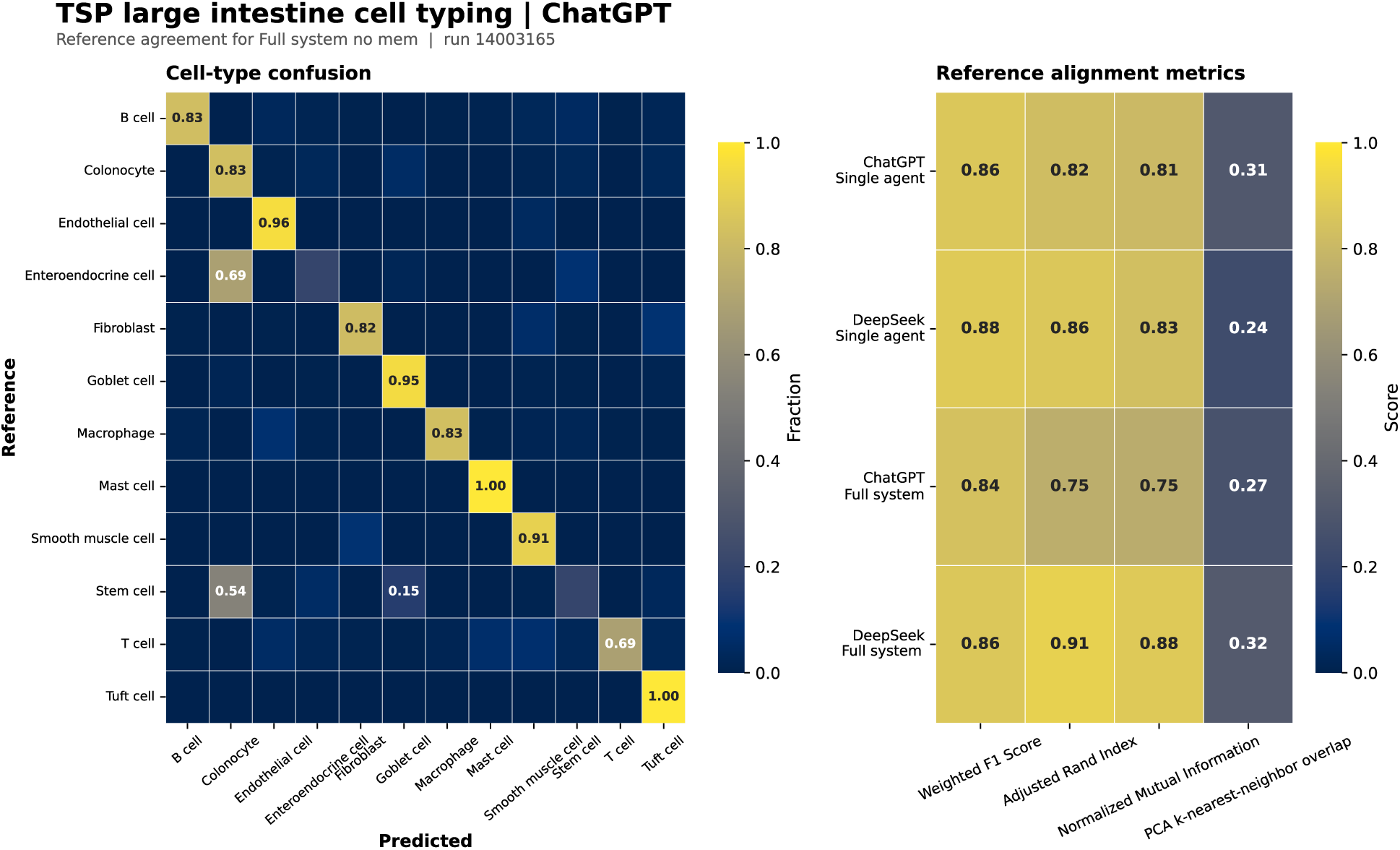
Cell Type Annotation Accuracy, Tabula Sapiens Large Intestine. (*Left*) Cell-type confusion matrix for the ChatGPT Full Agent System run on the Tabula Sapiens Large Intestine dataset, showing predicted versus reference cell type assignments for 12 populations. (*Right*) Reference alignment metrics heatmap comparing four system configurations (ChatGPT Single Agent, DeepSeek Single Agent, ChatGPT Full System, DeepSeek Full System) across Weighted F1 Score, Adjusted Rand Index (ARI), Normalized Mutual Information (NMI), and PCA k-nearest-neighbor overlap.

**Figure 8.**
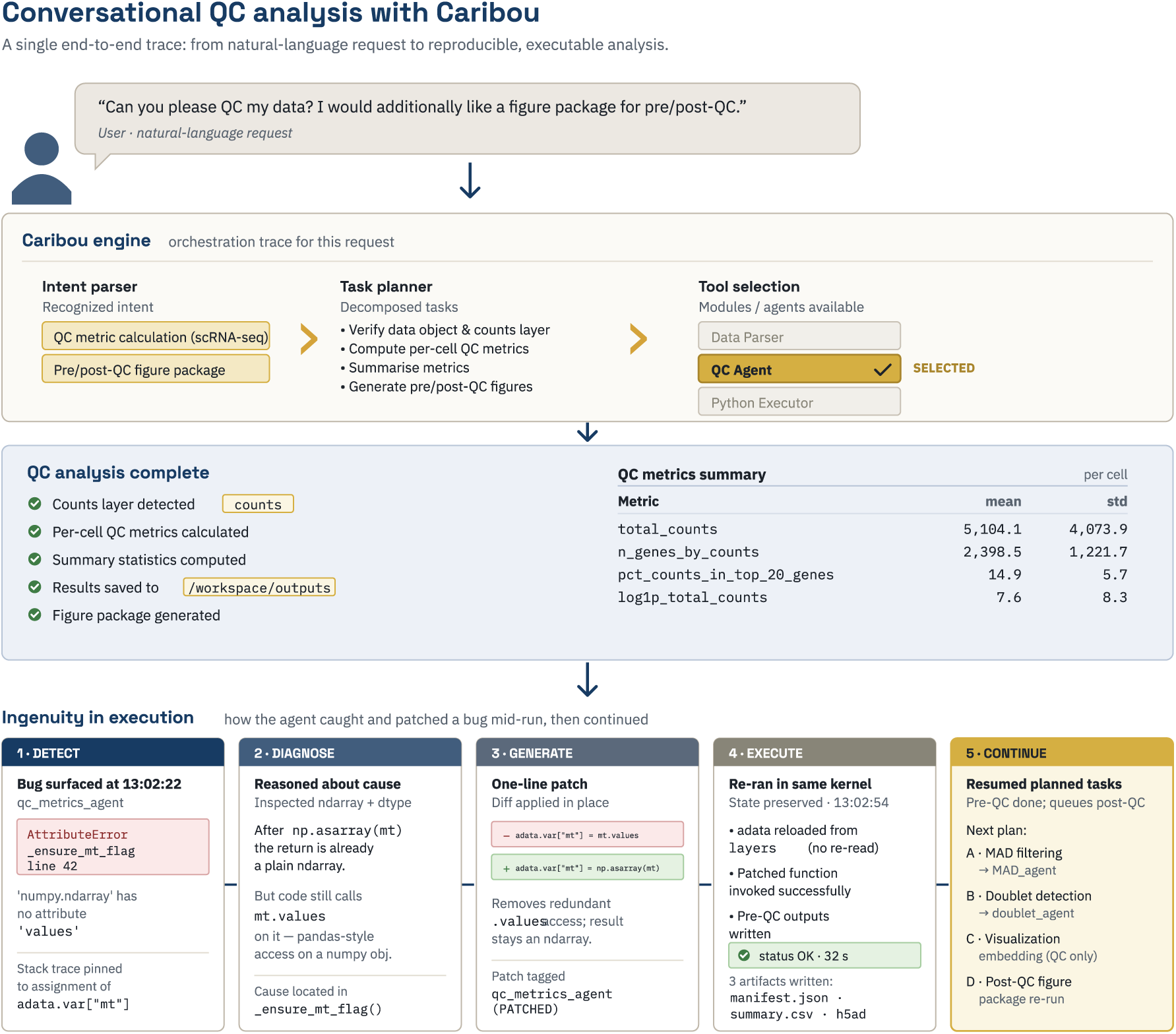
Adaptive Problem-Solving Case Studies. (*Case A*) Detailed trace of an adaptive error-recovery case: CARIBOU is asked to plot specific QC metrics, detects that required fields are absent from the AnnData object, autonomously generates and executes the code to compute them, then proceeds with the original plotting request.

The repository includes a conventional software test suite covering the agent-system loader, message parsing, memory manager behavior, API client wrappers, and end-to-end message flow, providing validation at the orchestration level independent of biological benchmarks.

### Accessibility

We chose to use the colormap “cividis” as the default option for generating figure output within CARIBOU as well as for the figures included in this manuscript in order to make images accessible to individuals with color vision deficiency (CVD). Cividis is optimized for similar visual interpretation between those with and without CVD while maintaining hue and brightness (Nuñez et al., 2018).

## RESOURCE AVAILABILITY

### Lead contact

Requests for additional information or resources should be directed to the lead contact, Dr. Matthew Rose, who will ensure that these are provided.

### Materials availability

This study did not generate new materials.

### Data and code availability

The datasets used in this paper are publicly available from the sources described in the Methods. All source code generated during this study has been deposited in GitHub.

https://github.com/rose-research-group/CARIBOU

## ACKNOWLEDGEMENTS

We are indebted to the broader open-source and computational biology communities whose tools, datasets, and infrastructure made this work possible. We thank Drs. Barbara Jusiak and Susan Rose for their invaluable discussions and feedback throughout development of the project. We also thank the maintainers and contributors of Scanpy, scVI-tools, HarmonyPy, Scrublet, CellTypist, and the AnnData ecosystem for providing foundational software infrastructure for single-cell analysis. We thank the Allen Institute for Brain Science and the Tabula Sapiens Consortium for generating and openly releasing the datasets used in this work, as well as the Gene Expression Omnibus (GEO) and CZI CELLxGENE platforms for supporting open biological data access.

## AUTHOR CONTRIBUTIONS

D.R. and N.S. jointly conceived and developed the CARIBOU framework. D.R. led implementation of the core framework architecture, execution environment, and benchmarking infrastructure. N.S. led development of the agent systems, workflow blueprints, orchestration logic, and prompting strategies. D.R. and N.S. jointly designed benchmarking experiments, evaluated system behavior, analyzed results, and contributed substantially to writing and revising the manuscript.

P.S. contributed to software development, testing, workflow evaluation, and benchmarking support. V.V. contributed to software development, benchmarking, testing, and figure review. M.F.R. supervised the study, contributed to project direction and conceptual framing, and edited the manuscript.

## SUPPLEMENTAL INFORMATION

**Figure S1.**
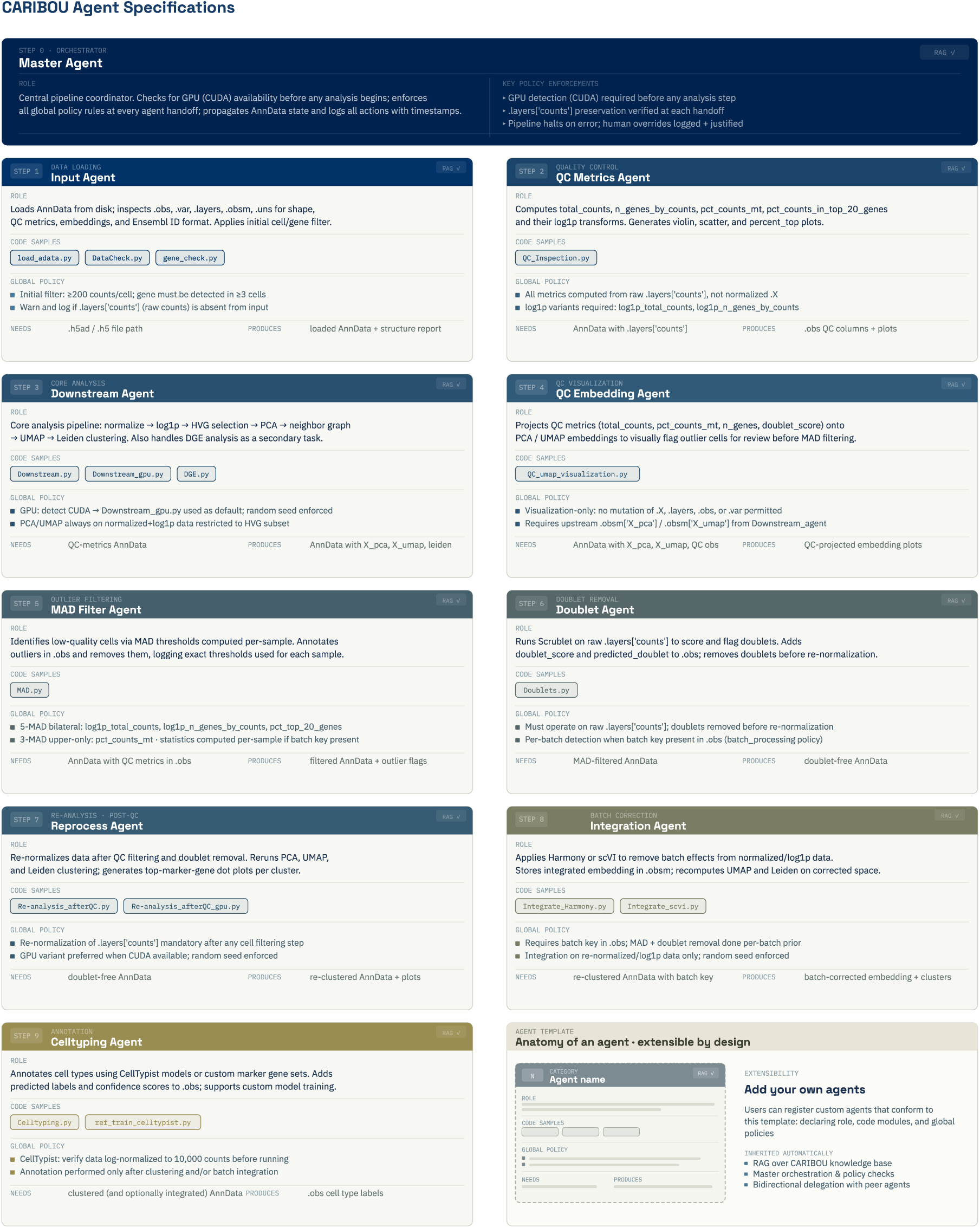
CARIBOU Agent Specification Cards. Each card describes one specialist agent in the fully connected blueprint, including its role, representative code samples, global policy constraints, required inputs, and produced outputs.

**Figure S2.**
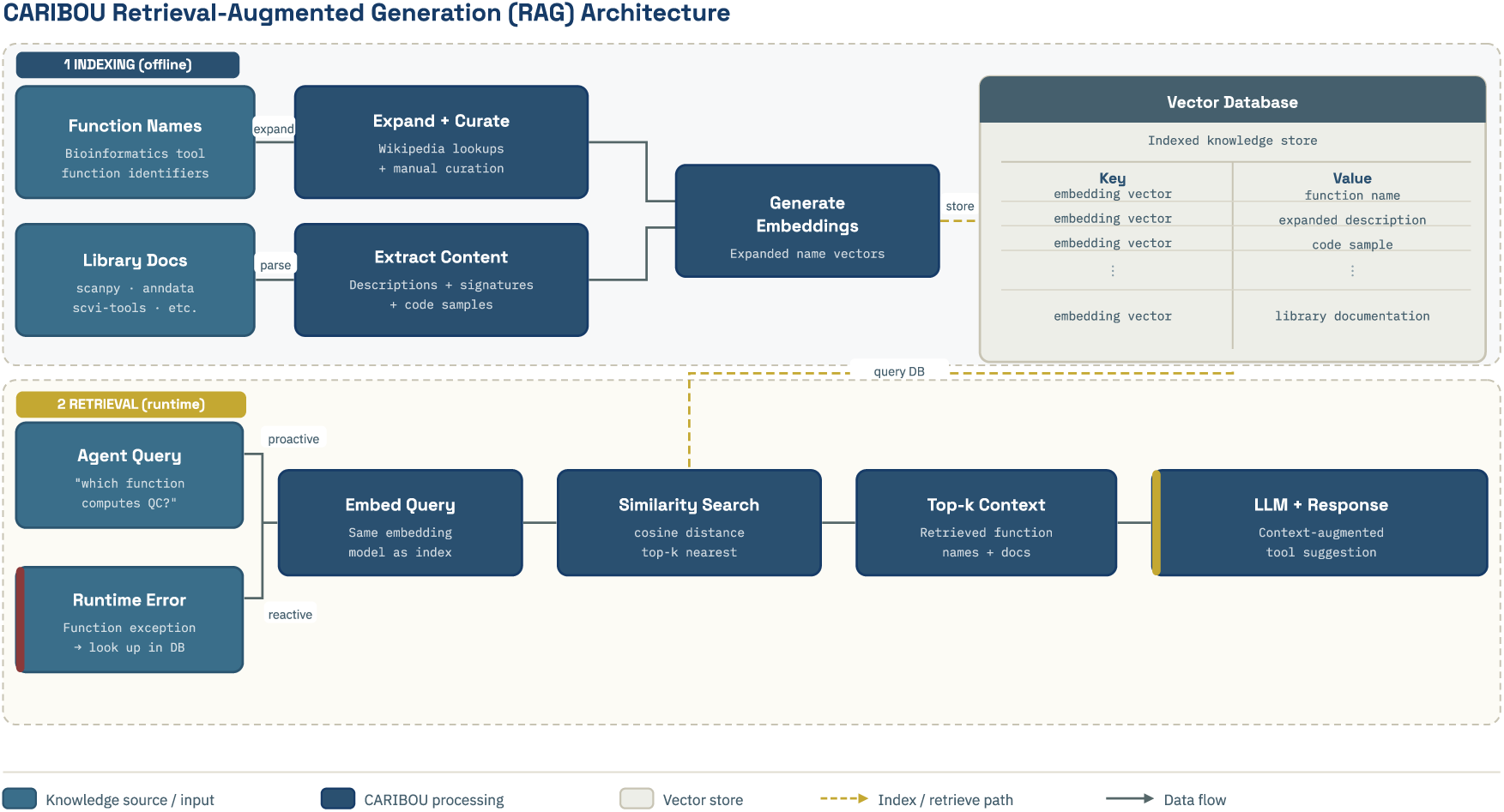
CARIBOU Retrieval-Augmented Generation (RAG) Architecture. The RAG module operates in two phases. During offline indexing, bioinformatics function names and library documentation are expanded, curated, and embedded using a local embedding model, then stored in a vector database keyed by embedding vector with associated function names, descriptions, code samples, and documentation. During runtime retrieval, agent queries and runtime execution errors trigger embedding-based similarity search over the vector store, returning top-k context entries that are injected into the active agent context to guide code generation and error correction.

**Figure S3.**
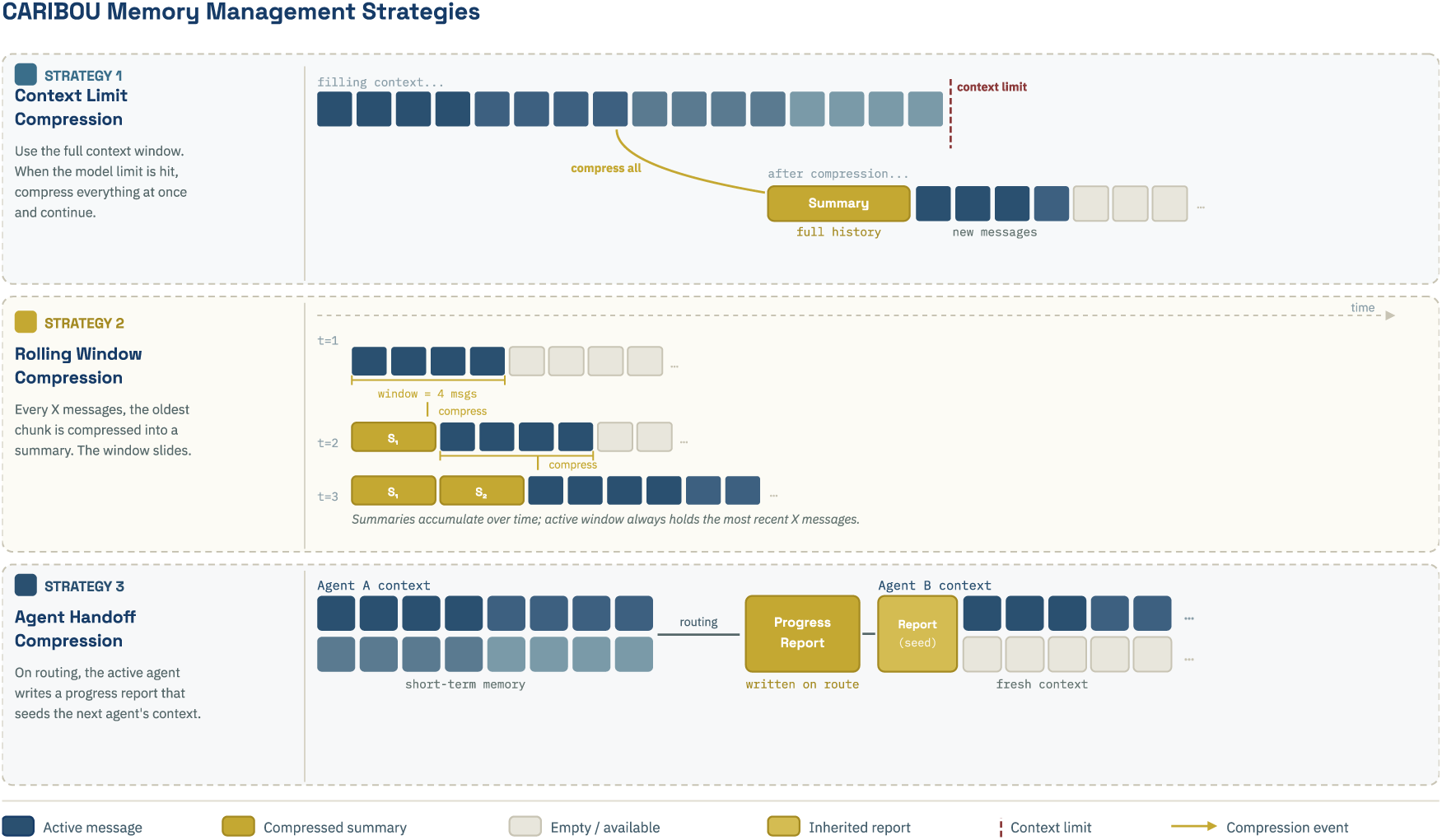
CARIBOU Context Management Strategies. Three strategies for managing conversational history across long multi-agent sessions are illustrated. Strategy 1, context limit compression, maintains the full growing transcript until the model context limit is approached, at which point all history is compressed into a single summary and the session continues. Strategy 2, rolling window compression, periodically compresses the oldest message chunks into summaries while retaining a recent uncompressed tail, allowing the active window to always hold the most recent interactions. Strategy 3, agent handoff compression, generates a structured progress report at each delegation event, which seeds the incoming agent’s context in place of the full prior conversation history.

**Figure S4.**
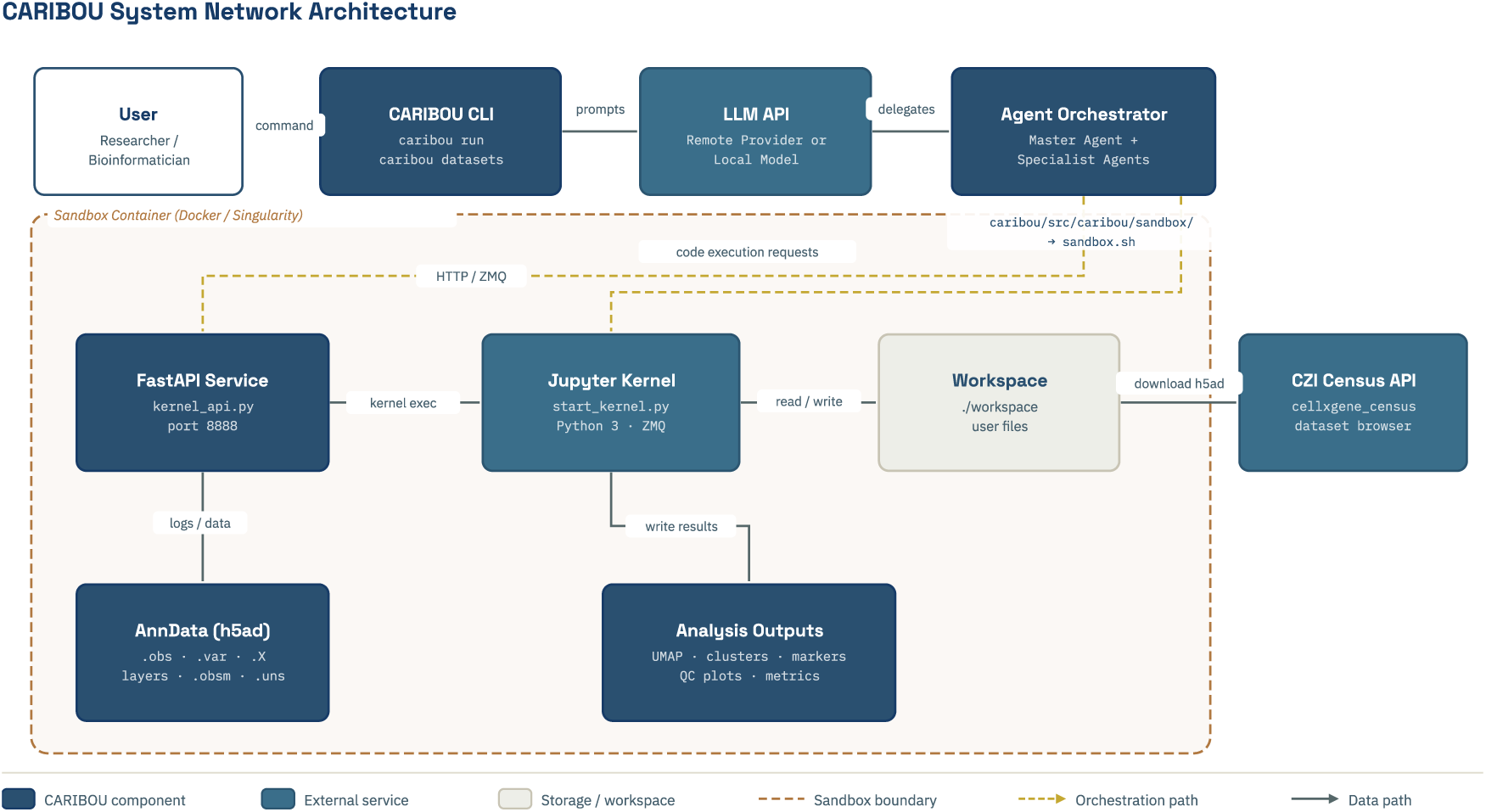
CARIBOU System Network Architecture. End-to-end data and orchestration flow from user command to analysis output. The user interacts with the CARIBOU CLI, which routes prompts to a remote or local LLM API and delegates execution to an agent orchestrator managing the Master Agent and specialist agents. Within the sandbox container boundary, a FastAPI service interfaces with a persistent Jupyter kernel for code execution. The kernel reads and writes to a shared workspace containing the AnnData object and analysis outputs. Input datasets can optionally be sourced from the CZI CELLxGENE Census API. Dashed lines indicate orchestration paths and sandbox boundaries; solid lines indicate data flow.

**Figure S5.**
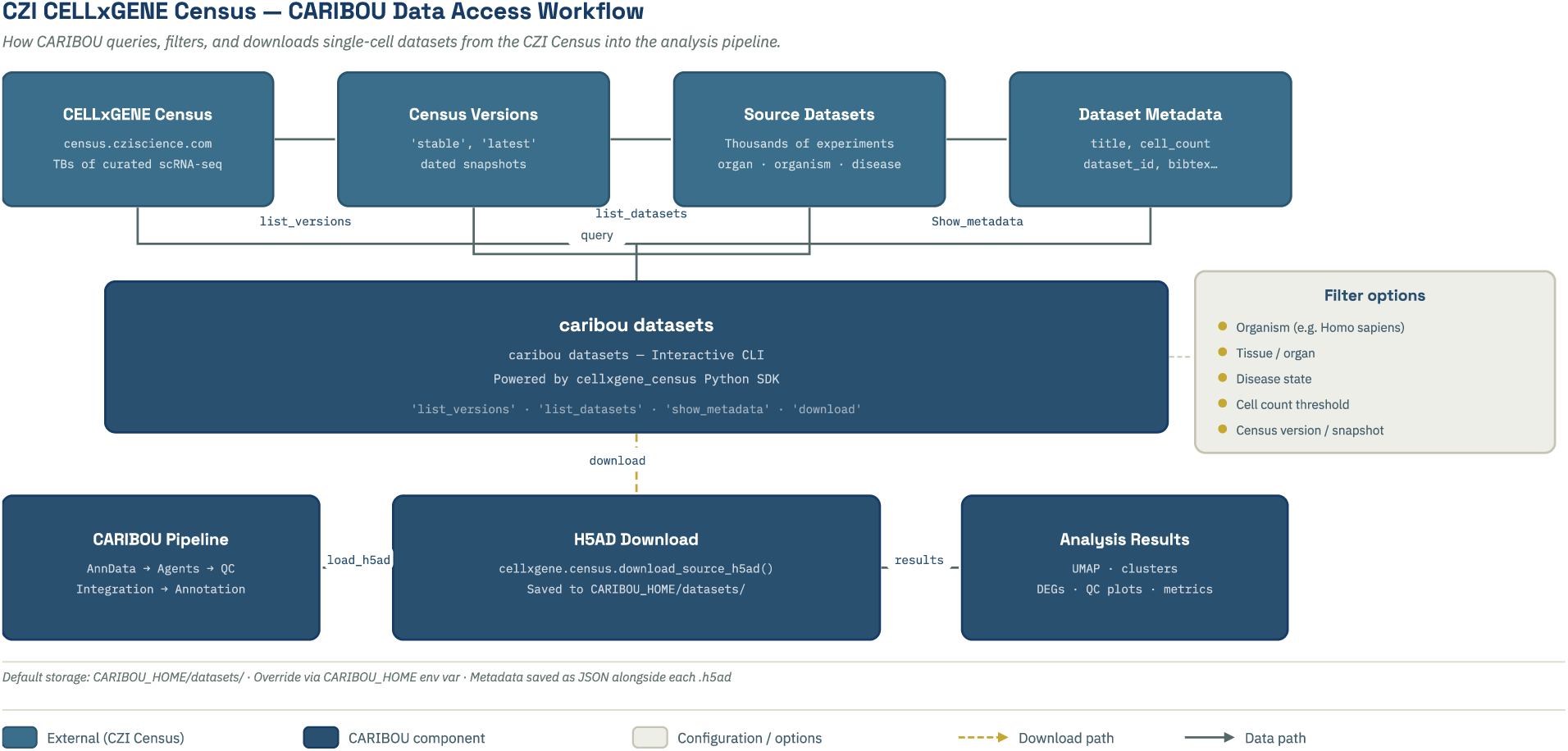
CZI CELLxGENE Census - CARIBOU Data Access Workflow. Schematic of how CARIBOU queries, filters, and downloads single-cell datasets from the CZI CELLxGENE Census into the analysis pipeline. The caribou datasets CLI command interfaces with the Census Python SDK to list available census versions and source datasets, display metadata, and apply filters including organism, tissue or organ, disease state, cell count threshold, and census version snapshot. Selected datasets are downloaded as AnnData h5ad files to local storage alongside a JSON metadata record, and are then passed directly into the CARIBOU analysis pipeline for QC, integration, and annotation.

**Figure S6.**
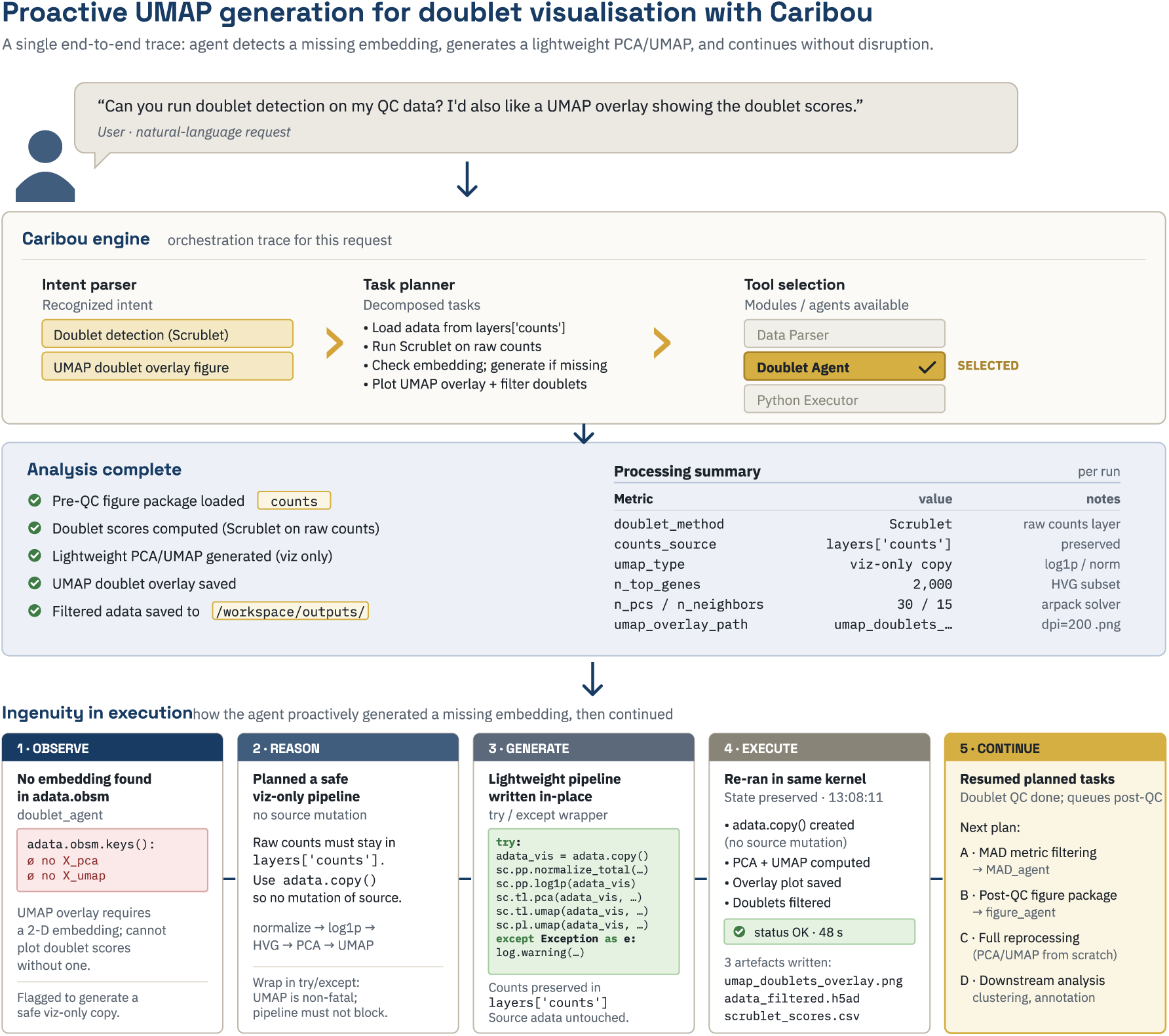
Adaptive Problem-Solving Case Study: Proactive UMAP Generation for Doublet Visualization. A single end-to-end execution trace illustrating proactive adaptive behavior. The user requests doublet detection with a UMAP overlay of doublet scores. Upon loading the data, the Doublet Agent observes that no embedding exists in the AnnData object. Rather than failing, the agent reasons that UMAP generation is non-fatal and plans a safe visualization-only pipeline using a copy of the data to avoid source mutation. The agent generates a lightweight normalization, PCA, and UMAP pipeline, executes it in the live kernel, produces the doublet overlay figure, and resumes the planned downstream task queue without disruption.

**Figure S7.**
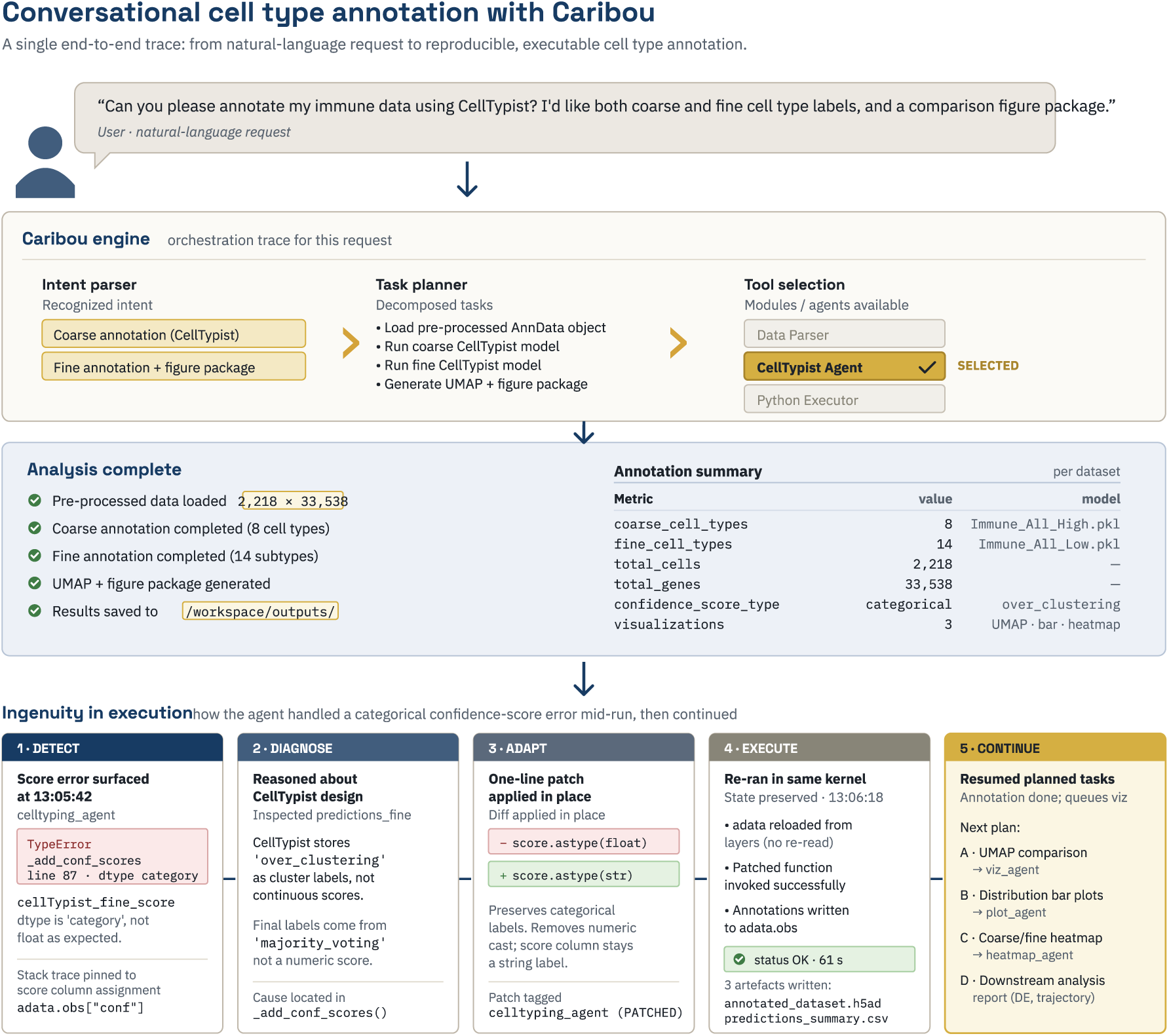
Adaptive Problem-Solving Case Study: Conversational Cell-Type Annotation with Mid-Run Error Recovery. A single end-to-end execution trace illustrating reactive error correction during CellTypist-based annotation. The user requests coarse and fine cell-type annotation with a comparison figure package. Mid-execution, the Celltyping Agent encounters a TypeError when attempting to cast CellTypist confidence scores from categorical to float, arising because CellTypist stores majority-voting labels as cluster labels rather than continuous scores. The agent diagnoses the cause, applies a one-line in-place patch converting the score column to string type, re-executes successfully in the same kernel with state preserved, writes annotated outputs to the workspace, and queues downstream visualization tasks.

