## Supplementary Figures for "CARIBOU: Computational AI Research Interface for Bioinformatics, Omics, and Unifying Agents"

### SUPPLEMENTAL INFORMATION

#### Supplemental Table of Contents


Supplemental Figure S1. CARIBOU Agent Specification Cards 35
Supplemental Figure S2. CARIBOU Retrieval-Augmented Generation Architecture 36
Supplemental Figure S3. CARIBOU Context Management Strategies 37
Supplemental Figure S4. CARIBOU System Network Architecture 38
Supplemental Figure S5. CZI CELLxGENE Census–CARIBOU Data Access Workflow 39
Supplemental Figure S6. Proactive UMAP Generation for Doublet Visualization 40
Supplemental Figure S7. Conversational Cell-Type Annotation with Mid-Run Error Recovery 41

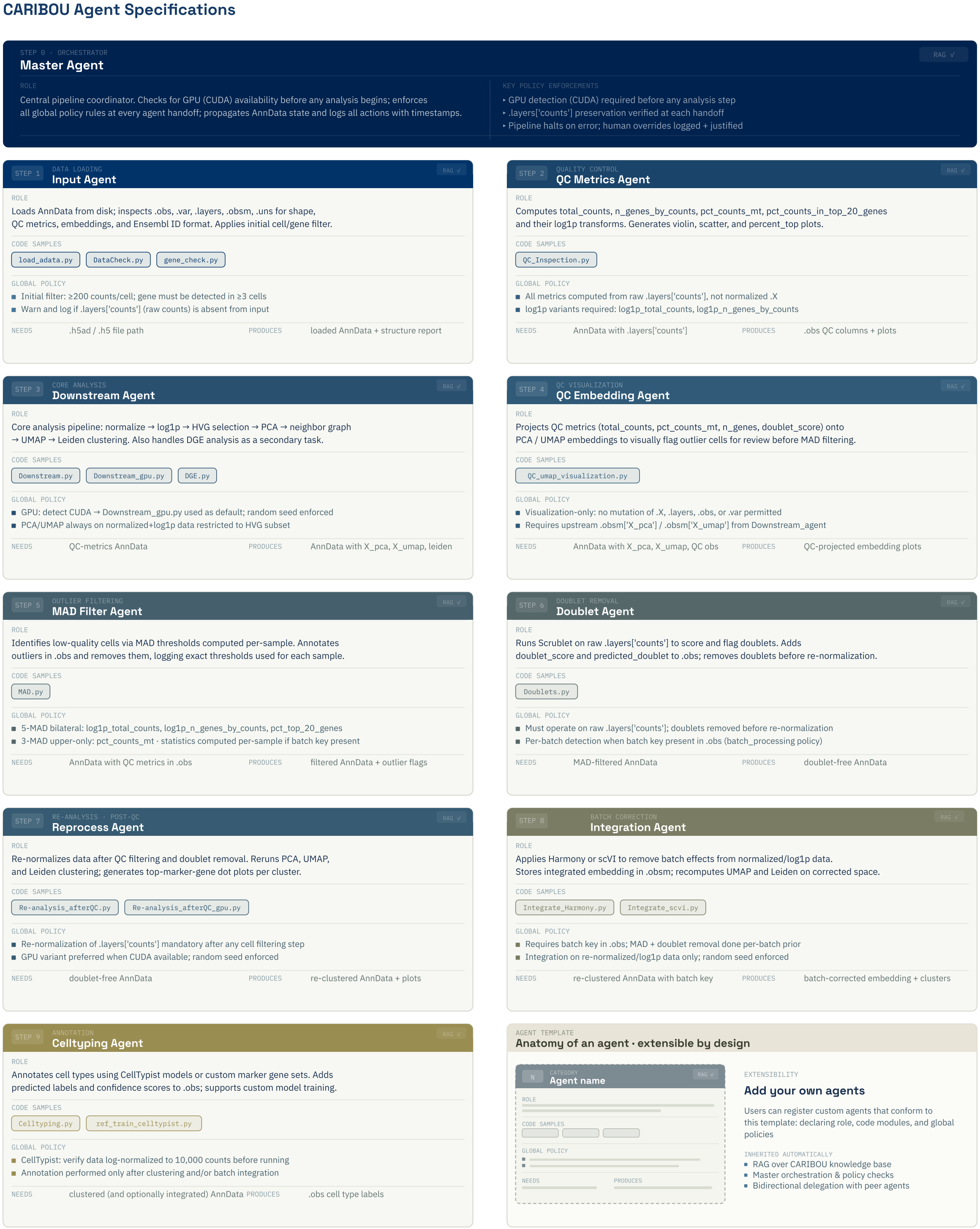


**Figure S1.** CARIBOU Agent Specification Cards. Each card describes one specialist agent in the fully connected blueprint, including its role, representative code samples, global policy constraints, required inputs, and produced outputs.

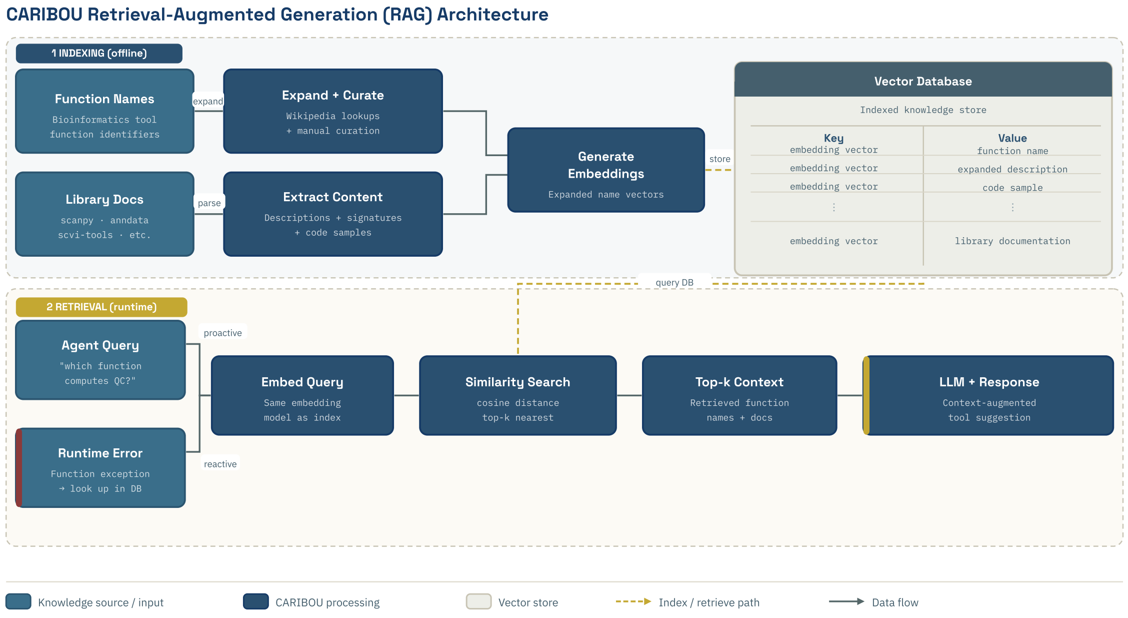


**Figure S2.** CARIBOU Retrieval-Augmented Generation (RAG) Architecture. The RAG module operates in two phases. During offline indexing, bioinformatics function names and library documentation are expanded, curated, and embedded using a local embedding model, then stored in a vector database keyed by embedding vector with associated function names, descriptions, code samples, and documentation. During runtime retrieval, agent queries and runtime execution errors trigger embedding-based similarity search over the vector store, returning top-k context entries that are injected into the active agent context to guide code generation and error correction.

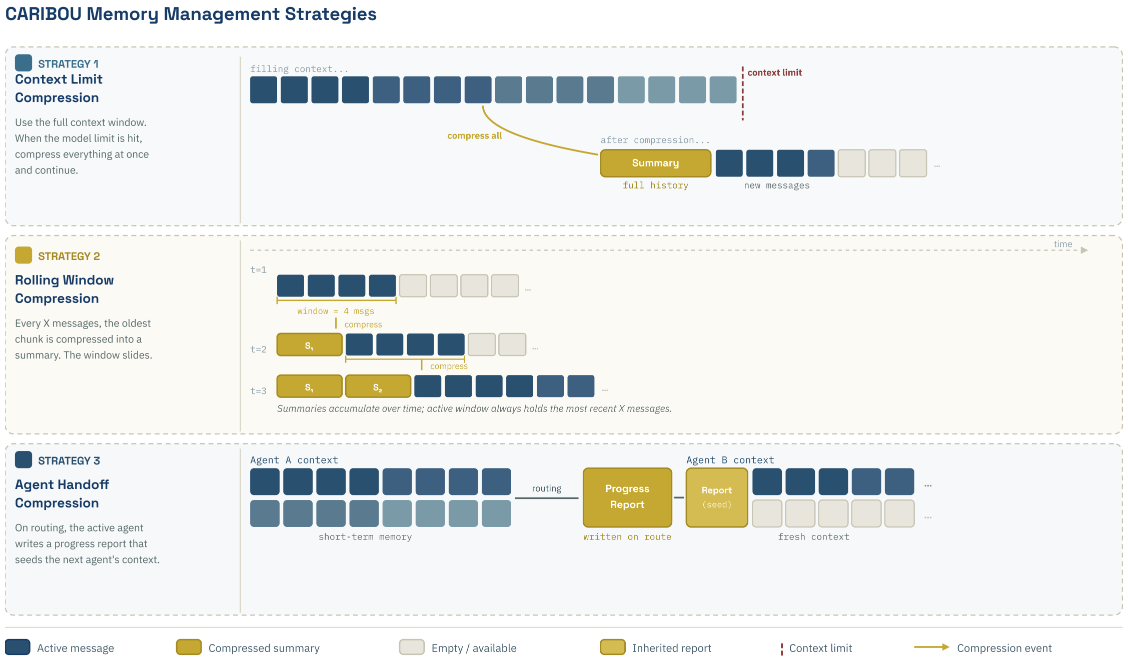


**Figure S3.** CARIBOU Context Management Strategies. Three strategies for managing conversational history across long multi-agent sessions are illustrated. Strategy 1, context limit compression, maintains the full growing transcript until the model context limit is approached, at which point all history is compressed into a single summary and the session continues. Strategy 2, rolling window compression, periodically compresses the oldest message chunks into summaries while retaining a recent uncompressed tail, allowing the active window to always hold the most recent interactions. Strategy 3, agent handoff compression, generates a structured progress report at each delegation event, which seeds the incoming agent's context in place of the full prior conversation history.

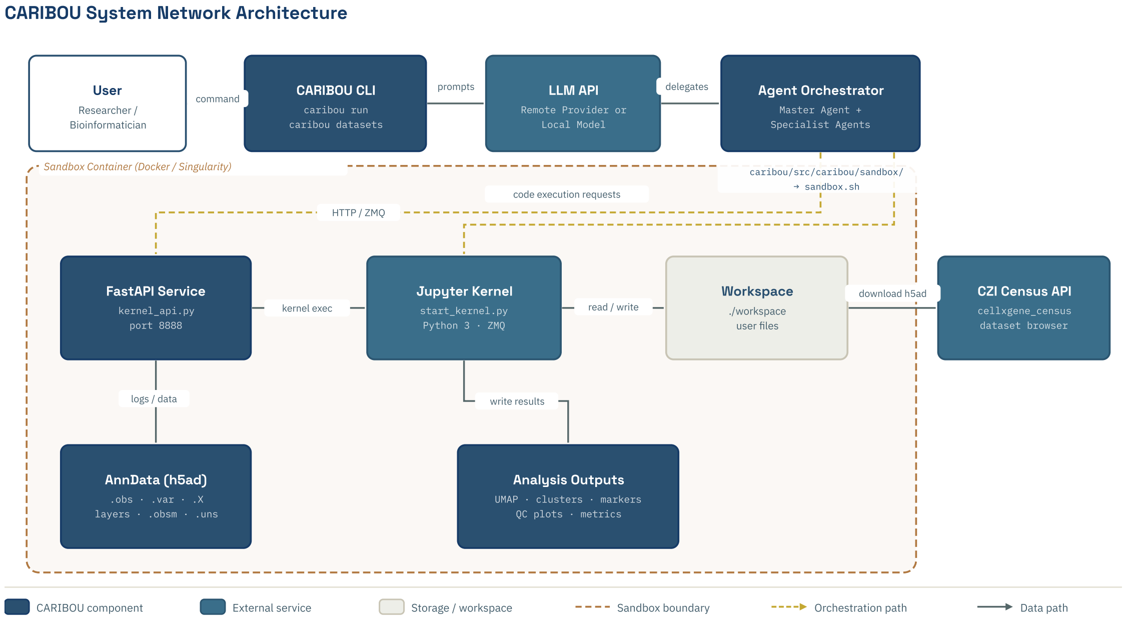


**Figure S4.** CARIBOU System Network Architecture. End-to-end data and orchestration flow from user command to analysis output. The user interacts with the CARIBOU CLI, which routes prompts to a remote or local LLM API and delegates execution to an agent orchestrator managing the Master Agent and specialist agents. Within the sandbox container boundary, a FastAPI service interfaces with a persistent Jupyter kernel for code execution. The kernel reads and writes to a shared workspace containing the AnnData object and analysis outputs. Input datasets can optionally be sourced from the CZI CELLxGENE Census API. Dashed lines indicate orchestration paths and sandbox boundaries; solid lines indicate data flow.

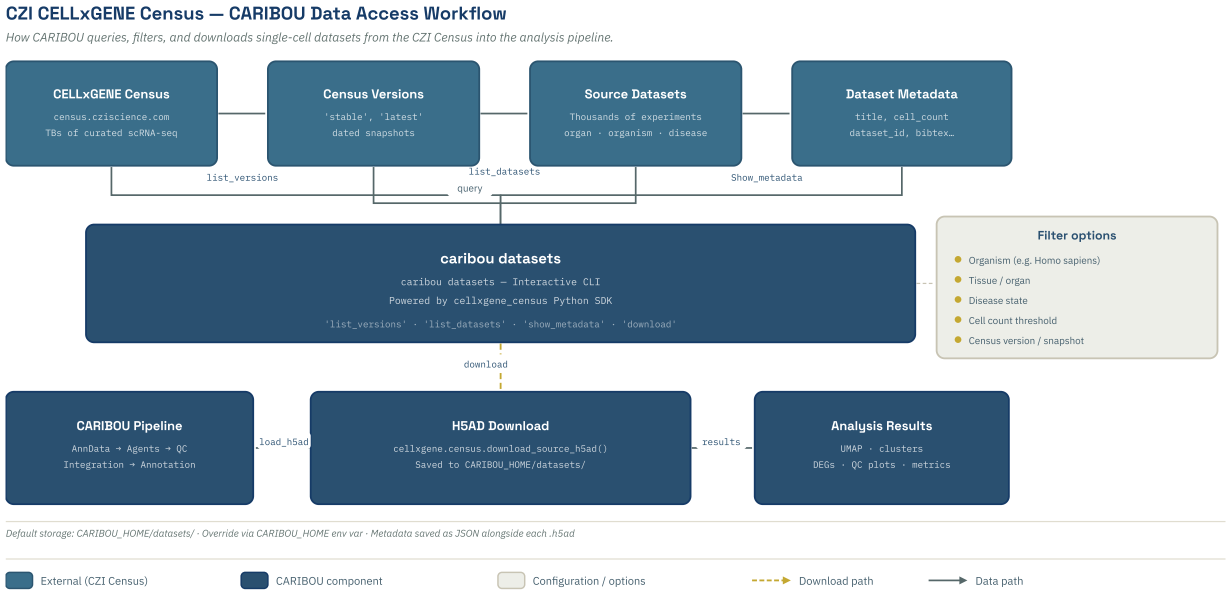


**Figure S5.** CZI CELLxGENE Census - CARIBOU Data Access Workflow. Schematic of how CARIBOU queries, filters, and downloads single-cell datasets from the CZI CELLxGENE Census into the analysis pipeline. The caribou datasets CLI command interfaces with the Census Python SDK to list available census versions and source datasets, display metadata, and apply filters including organism, tissue or organ, disease state, cell count threshold, and census version snapshot. Selected datasets are downloaded as AnnData h5ad files to local storage alongside a JSON metadata record, and are then passed directly into the CARIBOU analysis pipeline for QC, integration, and annotation.

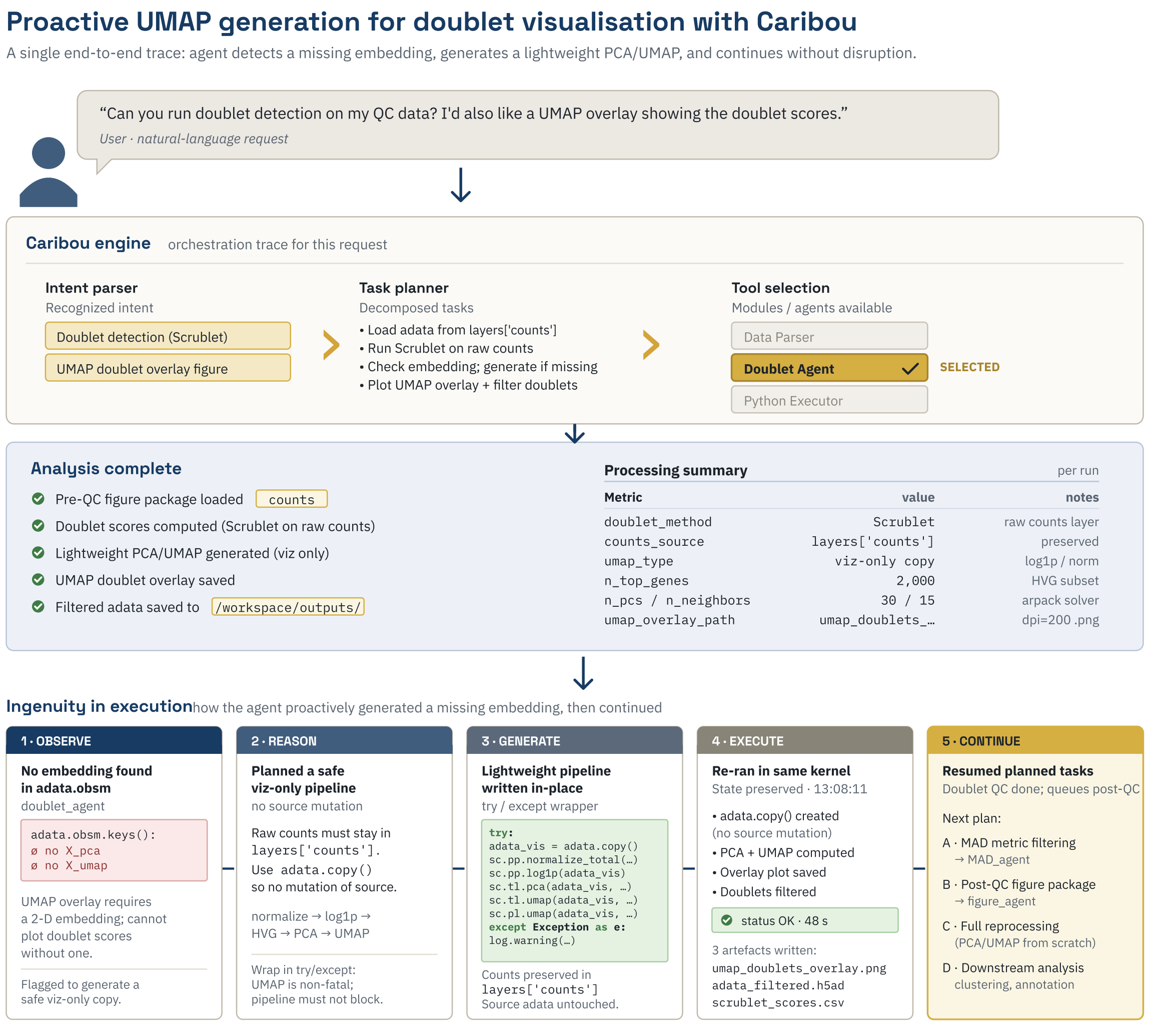


**Figure S6.** Adaptive Problem-Solving Case Study: Proactive UMAP Generation for Doublet Visualization. A single end-to-end execution trace illustrating proactive adaptive behavior. The user requests doublet detection with a UMAP overlay of doublet scores. Upon loading the data, the Doublet Agent observes that no embedding exists in the AnnData object. Rather than failing, the agent reasons that UMAP generation is non-fatal and plans a safe visualization-only pipeline using a copy of the data to avoid source mutation. The agent generates a lightweight normalization, PCA, and UMAP pipeline, executes it in the live kernel, produces the doublet overlay figure, and resumes the planned downstream task queue without disruption.

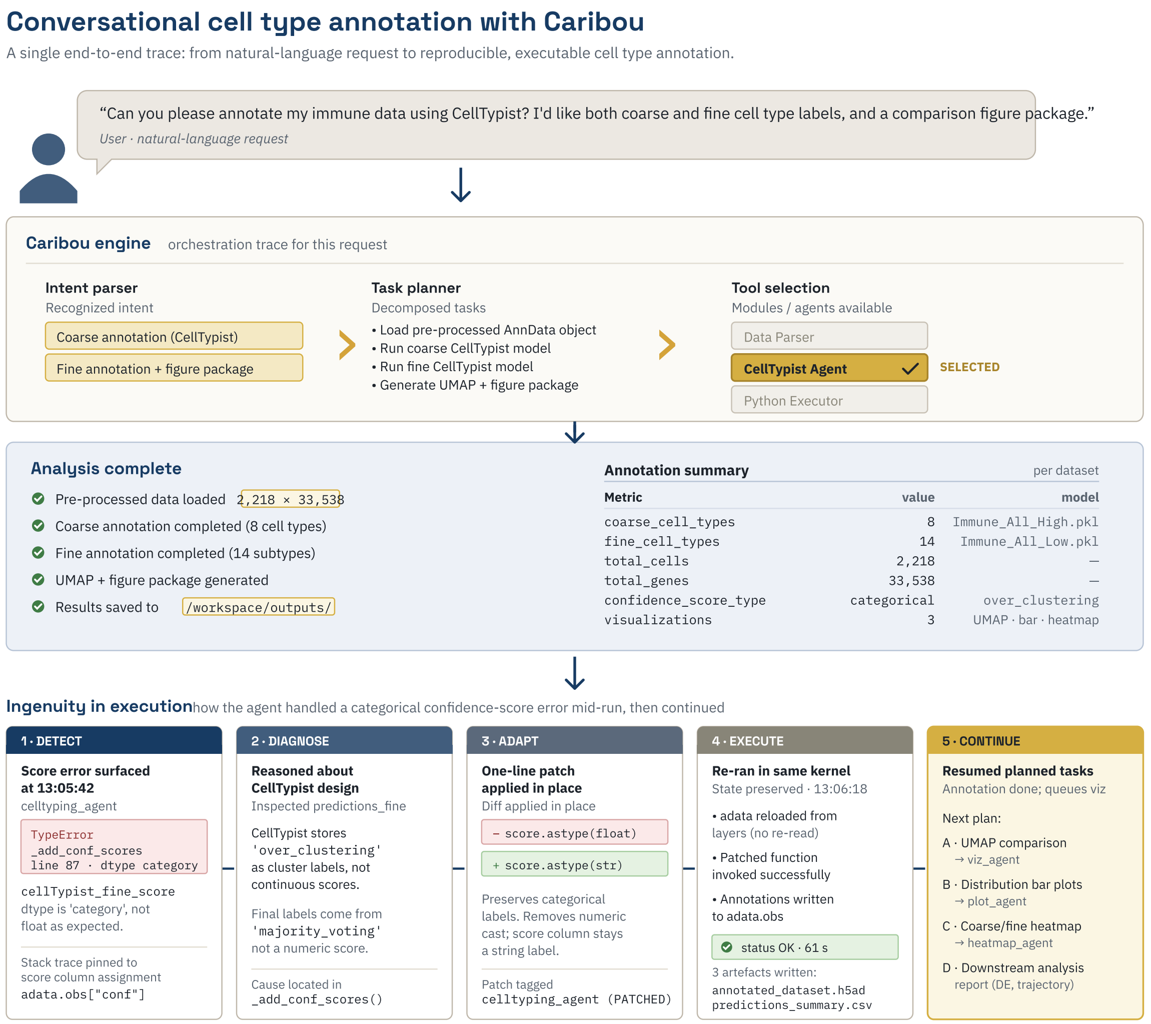


**Figure S7.** Adaptive Problem-Solving Case Study: Conversational Cell-Type Annotation with Mid-Run Error Recovery. A single end-to-end execution trace illustrating reactive error correction during CellTypist-based annotation. The user requests coarse and fine cell-type annotation with a comparison figure package. Mid-execution, the Celltyping Agent encounters a TypeError when attempting to cast CellTypist confidence scores from categorical to float, arising because CellTypist stores majority-voting labels as cluster labels rather than continuous scores. The agent diagnoses the cause, applies a one-line in-place patch converting the score column to string type, re-executes successfully in the same kernel with state preserved, writes annotated outputs to the workspace, and queues downstream visualization tasks.
